# Global quantitative understanding of nonequilibrium cell fate decision making in response to pheromone

**DOI:** 10.1101/2022.07.04.498707

**Authors:** Sheng Li, Qiong Liu, Erkang Wang, Jin Wang

## Abstract

Cell cycle arrest and polarized cell growth are commonly used to qualitatively characterize the fate of yeast in response to pheromone. However, the quantitative decision-making process underlying the time-dependent changes in cell fate remains unclear. Here, by observing the multi-dimensional responses at the single-cell level experimentally, we find that yeast cells have various fates. Multiple states are revealed, along with the kinetic switching rates and pathways among them, giving rise to a quantitative landscape of mating response. We developed a theoretical framework using a nonequilibrium landscape and flux theory to account for the cell morphology observed experimentally and performed a stochastic simulation of biochemical reactions to explain the signal transduction and cell growth. Our experimental results established the first global quantitative demonstration of the real-time synchronization of intracellular signaling with their physiological growth and morphological functions which reveals the underlying physical mechanism. This study provides an emerging mechanistic approach for understanding the nonequilibrium global pheromone-regulated cell fate decision-making in growth and morphology.

## INTRODUCTION

Facing a variety of external stimuli, cells integrate information from various sources and initiate appropriate stress responses, such as survival or death, division or differentiation, and shape change or migration (*1–7*). To better understand the cellular response to external stimuli, we chose the pheromone pathway of *Saccharomyces cerevisiae*, which is a classical model of the mitogen-activated protein kinase (MAPK) signaling pathway (*8–11*). *Saccharomyces cerevisiae* has two modes of reproduction, sexual mating reproduction and asexual budding reproduction (*12–16*). These two reproductive modes can be converted either with or without the assistance of external pheromone stimulation from a partner (*17*). For sexual reproduction, mating to form a diploid cell is the natural behavior of heterosexual haploid *Saccharomyces cerevisiae* to cope with an unfavorable environment and to improve their survival rate for generations (*18–20*). Pheromones, a type of sex hormone secreted by *Saccharomyces cerevisiae*, serve as the yeast’s mating signal, informing its partner to prepare for the beginning of cell fusion. (*18–26*). What will happen to the yeast cell if its partner does not arrive and this normal stimulation continues? From microscopic gene network **(Fig. 1)**, external signals are transmitted to Fus3 through a series of internal proteins such as prototype heterotrimeric GTP binding protein and the MAPK kinase cascade (*27–29*). Then, Fus3 shuttles back and forth across the nuclear membrane, directly or indirectly activating genes for cell cycle arrest (Far1) and polar growth (Bni1) (*30–38*). Thus, the current understanding of the pheromone-induced fate of yeast cells can be qualitatively summarized as the occurrence of cell cycle arrest and polarized cell growth (*39–41*).

**Fig. 1.**
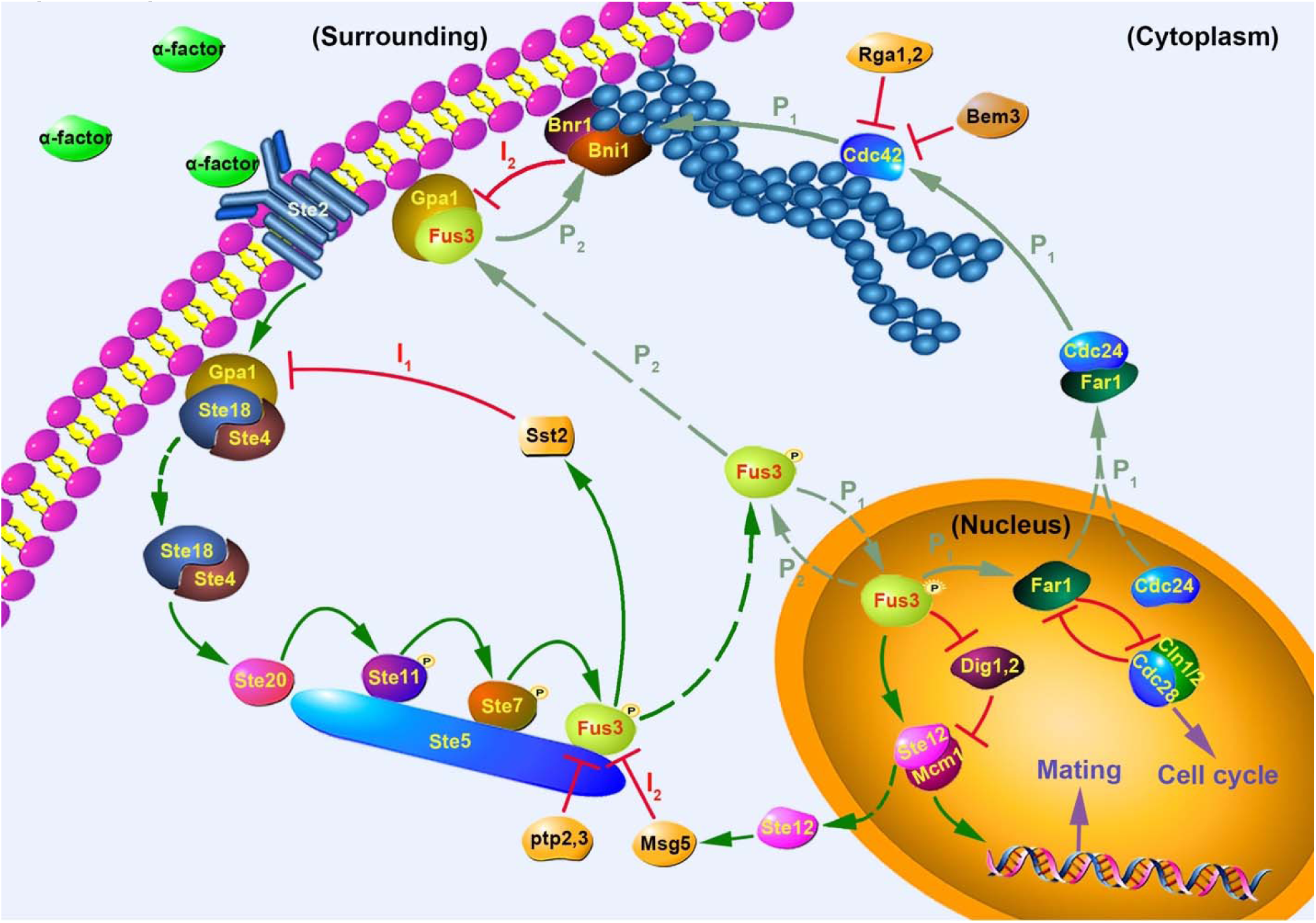
Schematic diagram of the mating signal pathway of the yeast cell pheromone pathway. The red horizontal line represents the inhibition of negative feedback; the green arrow represents the activation of positive feedback; the dashed arrow represents a shift in localization; I_1_ and I_2_ represent different negative feedback adjustment pathways; and P_1_ and P_2_ represent the two polar growth signaling pathways that are connected by light green arrows.

During qualitative studies of cell fate, researchers have recognized that cellular decision-making plays a pivotal role in determining the ultimate fate of cells in the presence of various external stimuli (*42–54*). From a biological perspective, cell fate decisions can be conceptually characterized as the cascading transmission of signals along a static causal pathway (*55–59*). However, the quantitative dynamics of cell fate decision-making over time during global responses at both the mesoscopic and microscopic levels remain poorly understood. Recent advances in live cell fluorescence imaging platforms have led to increasing focus on studying the dynamics of biological processes and the underlying molecular mechanisms in living cells (*54, 60–73*). Nevertheless, disruptions in the external environment can perturb the delicate balance of cellular activities and exacerbate internal imbalances in yeast cells, as living cells are complex non-equilibrium microsystems (*74–83*). Therefore, quantifying these underlying cell fate decisions in response to external stimuli remains a significant challenge.

Indeed, from a microscopic perspective, the cellular decision-making of yeast in response to mating information is accomplished through multiple feedback loops, not only including the positive feedback regulation mediated by Fus3, but also through certain negative feedback regulation that can effectively reduce the transcriptional output of this pathway (*84–89*). After pheromone stimulation, there are two pathways that can lead to polar growth of the cells (**Fig. 1**). The first path (P_1_) is “Fus3 → Far1/Cdc24 → Cdc42 → Bni1” (*40, 90–93*). This is realized by Fus3 entering the nucleus to activate Far1, which can escape from the nucleus to activate Bni1 indirectly. The second path (P_2_) is direct activation by Fus3 of Bni1 in the cytoplasm, “Fus3 → Bni1” (*31, 94*). In the process of these two polar growth pathways acting alone or in concert, the pheromone-induced self-activation of Fus3 (“Fus3 → Fus3”) facilitates the rapid transmission of the signals (*95–98*). Activation of the pheromone pathway also induces multiple negative feedback loops, such as I_1_ (Sst2 and Gpa1) (*87-89, 99-101*), I_2_ (Msg5 and Bni1) (*31, 85-89, 102-104*) in the pathway. Only at high-dose pheromone levels is the negative feedback of the downstream Msg5 upregulated (*85*) and only at high concentrations of Bni1 is activity of Fus3 indirectly reversed by Bni1 (*31, 94, 105-107*).

Here, by observing the multi-dimensional response at the single-cell level, we discovered the non-equilibrium steady states, which demonstrate the cell fates quantitatively, in response to different concentrations of pheromone. These steady states or cellular destinies include two gene expression levels, four growth rates, and four morphological fates. We quantified these responsive fates of the yeast cells in real time from various dimensions, including expression levels of Fus3 inside and outside of the nucleus, cell morphology, cell growth rate, and the stimulation concentrations. Multiple states, as well as switching kinetic rates and pathways among them, were revealed, giving rise to a quantitative landscape of the mating response. The applications of landscape and flux theory to this biological system enabled us to quantify the non-equilibrium dynamics of the yeast cell mating responses. Our results established a global and physical framework for understanding cell fate decision making and mating dynamics. Furthermore, we experimentally validated the proposed molecular mechanism for the formation of these states. These molecular mechanisms establish the links between molecular events to cellular characteristics across scales. They allow for the real-time synchronization of intracellular signaling with their physiological growth and morphological functions, bridging microscopic mechanisms and mesoscopic functions. To further elucidate these microscopic mechanisms, we conducted biochemical reaction simulations to demonstrate the emergence of these states. These findings shed new light on the global signaling mechanisms that govern how yeast determines its cellular fate in response to pheromone.

## RESULTS

### Quantifying the cellular decision-making threshold

Studies have shown that yeast, for which mating decisions are an all-or-none switch-like response, can automatically filter out very low-dose pheromone signals and only respond near a critical concentration or higher (*101, 108, 109*). This is due to the fact that very low doses of pheromone induce insufficient accumulation of intracellular signals, such as Fus3, which are incapable of initiating the mating-readiness project (*110–112*). To determine a pheromone concentration capable of arresting the cell cycle, we measured the fluorescence intensity of *FUS3*-GFP by flow cytometry in yeast cells incubated for 24 h with varying pheromone concentrations **(Fig. 2A)**. According to the fitted statistics, the expression of Fus3 at a pheromone dose of 0.01–0.4 μM showed unimodal distribution of the fluorescence intensities. The bimodal distribution representing the bistable states gradually began to appear when the dose was about 0.6 μM.

**Fig. 2.**
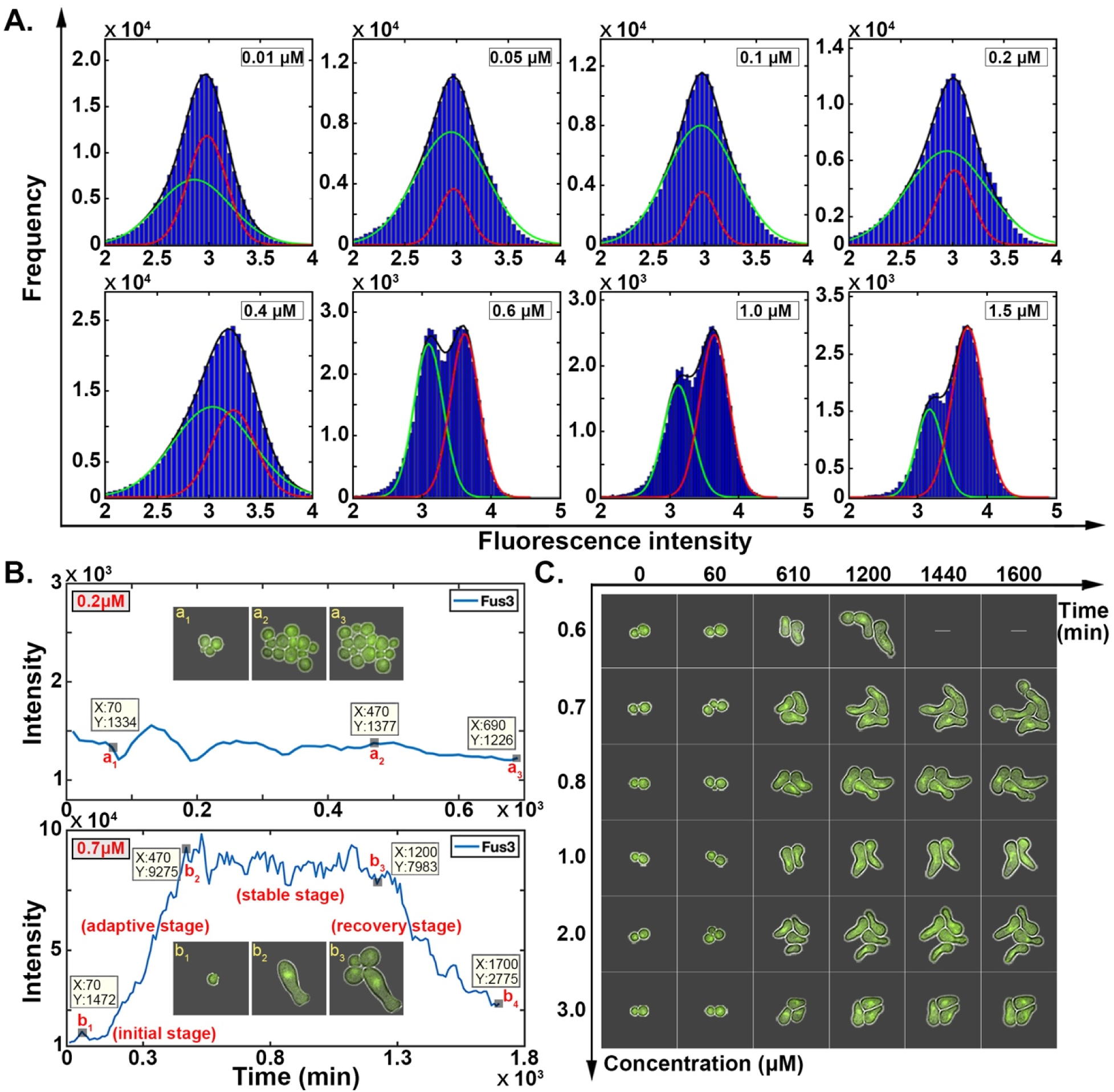
A non-equilibrium biological model for cellular responses. **(A)** The fluorescence intensity of *FUS3*-GFP was measured using flow cytometry. The yeast cells were cultured in YPD medium containing different concentrations of pheromone for 24 h; the black curve represents the overall fluorescence intensity statistics; and the green and red curves represent the two fitted statistical peaks. The sample size for each pheromone concentration test was 105,000 yeast cells. **(B)** Fus3 gene expression was observed microscopically in response to 0.2 μM and 0.7 μM pheromone. The x-axis represents the duration of exposure to the pheromone-containing medium. The y-axis represents the fluorescence intensity of *FUS3*-GFP; and images a_1_–a_3_ and b_1_–b_3_ depict the living states of yeast cells as observed through a microscope at their corresponding times. **(C)** The microscopically captured living state of yeast at various pheromone concentrations. The green fluorescence in cells represents the expression of *FUS3*-GFP; 60 min on the x-axis represents the time when yeast cells were switched to a culture medium containing pheromone.

In order to comprehend the correlation between gene expression levels of Fus3 and the threshold of cellular response, the fluorescence intensity trajectories of Fus3 expression in single cells were associated with the microscopy-based cellular response under various pheromone concentrations over time. A microfluidic device was used to culture yeast under constant temperature conditions in real time. Using total internal reflection fluorescence microscopy, the survival status and fluorescent signals of the yeast cells were captured in real time **(Movies S1– S7)**. Compared to 0.7 μM pheromone, the expression of Fus3 in the yeast cells did not fluctuate significantly at 0.2 μM, resulting in its inability to inhibit cell budding reproduction **(Fig. 2B)**. However, the expression of Fus3 at a pheromone concentration of 0.7 μM exhibited four stages: the initial stage (0–b_1_), the adaptive stage (b_1_–b_2_), the stable stage (b_2_–b_3_), and the recovery stage (b_3_–b_4_), corresponding to the biological behavior of yeast cells at different stages. Therefore, the unimodal distribution of 0.01–0.4 μM pheromone occurred when only the initial cells (0-b_1_) were present in the flow cytometer sample. This very low dose of pheromone was unable to arrest the cell cycle from the beginning in yeast. If the pheromone had arrested the cell cycle of yeast cells from the beginning, but not for more than 24 h, some yeast cells would have resumed budding at a later stage. As demonstrated by the bimodal distribution of 0.4–1.5 μM pheromone, the sample contained yeast cells in two distinct states (b_2_–b_3_ and b_3_–b_4_).

The results measured by flow cytometry were only rough estimates of critical mating decision thresholds, whereas microscopy of individual cells cultured in microfluidics revealed more precise cellular decisions. During culturing with varying concentrations of pheromone, we found that the time required for yeast cells to arrest their cell cycle was directly proportional to the pheromone concentration **(Fig. 2C)**. By extending the duration of the pheromone stimulation, the yeast cells gradually adapted to the external surroundings and resumed budding reproduction.

### Two cell fate decision states reflected in the expressions of Fus3

To explore the yeast cell fate decision-making during the stable stage (b_2_–b_3_ in **Fig.2B**), the fluorescence intensity trajectories of Fus3 inside and outside the nucleus were plotted over time **(Fig. 3A and Fig. S1–S2)**. The trajectories demonstrated that after approximately 600 min of yeast cell cultivation, Fus3 entered a stable stage where its fluorescence fluctuated around a certain value. In non-equilibrium statistical physics, the fluorescence intensity curve in this stable stage can be interpreted as a non-equilibrium steady state, whereas the rising phase of the first 600 min can be interpreted as a non-equilibrium non-steady state, or the relaxation process of the response. To visualize the cell fate of yeast at the level of gene expression, the three-dimensional statistical distribution of Fus3 (cytoplasmic and nuclear fluorescent signals) was plotted during the non-equilibrium steady-state phases **(Fig. 3B and Fig. S3–S5)**. Clearly, the landscape of Fus3 displayed two peaks, which correspond to the two cell fates that existed both inside and outside the nucleus.

**Fig. 3.**
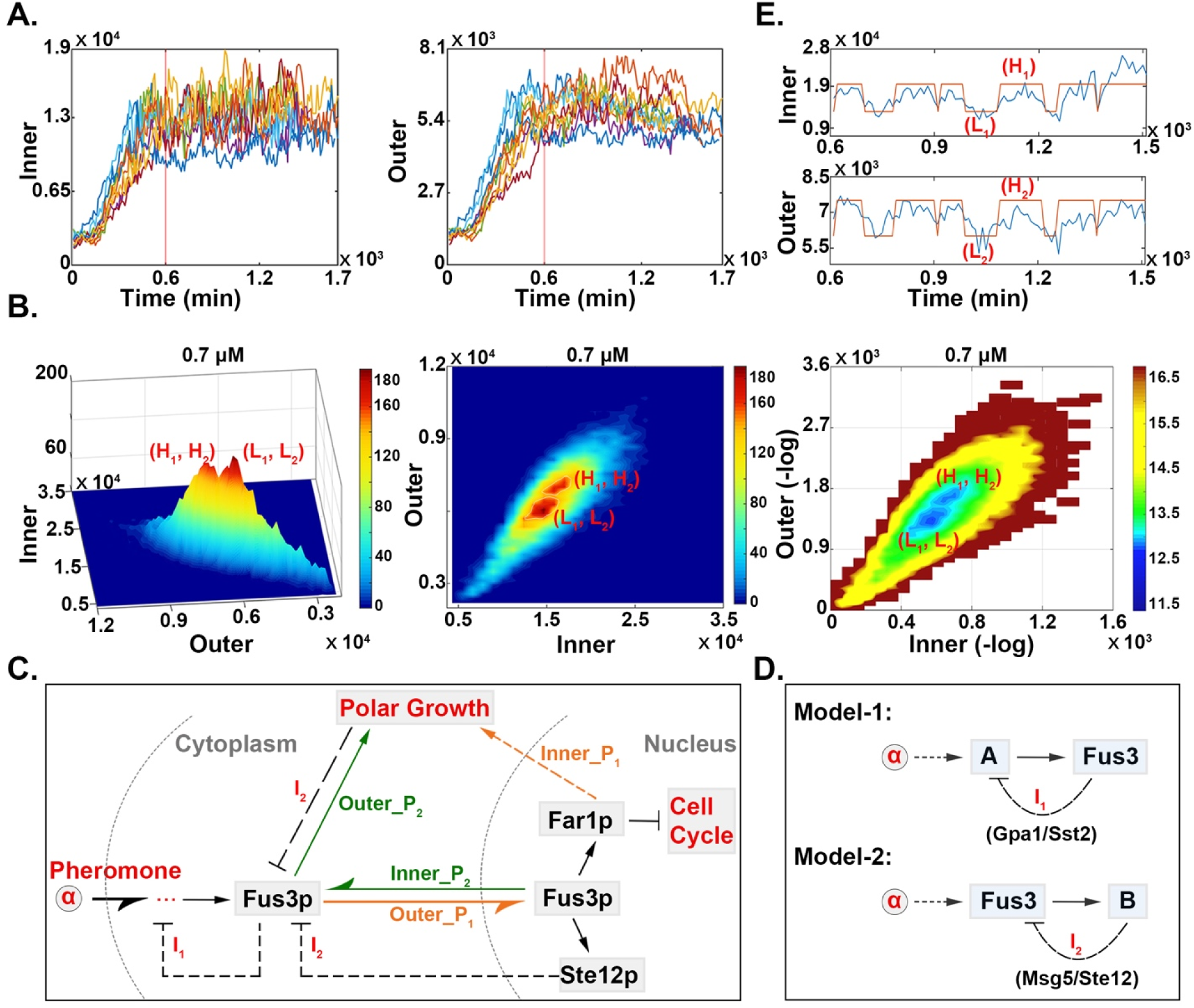
The two steady states of expression levels of Fus3. **(A)** Trajectories of fluorescence intensity of Fus3 at 0.7 μM inside and outside the nucleus (only a portion shown). The red vertical line at 600 min was used to approximate the time node at which all cell fluorescence trajectories had entered a non-equilibrium steady state. **(B)** Three-dimensional distribution graph of Fus3 fluorescence intensity inside and outside the nucleus of yeast cells in the stationary phase under different pheromone concentrations. On the left is the 3D distribution of fluorescence or the 3D population landscape, in the middle is the 2D histograms or the 2D underlying potential landscapes U in exponential scale (defined as p ∼ e^-U^), which is also the population landscape; on the right is the 2D underlying potential landscapes U (*U* = -lnP). The sample sizes at steady state for each pheromone concentration are as follows: 0.7 μM was equivalent to 21,335 cells, 0.8 μM to 18,408 cells, 1.0 μM to 23,041 cells, 2.0 μM to 36,886 cells, and 3.0 μM to 18,276 cells. **(C)** Diagram illustrating the molecular mechanism by which yeast cells respond to pheromone. The Outer_P_1_ represents the indirect pathway taken by Fus3 from the cytoplasm to the nucleus in order to inhibit the cell cycle; Inner_P_1_ represents the indirect pathway by which Fus3 in the nucleus promoted polar growth; Outer_P_2_ represents the direct pathway of Fus3 in the cytoplasm for polar growth; Inner_P_2_ represents the direct pathway for the transfer of Fus3 from the nucleus to the cytoplasm for polar growth; I_1_ andI_2_ represent the inhibitory effects of the negative feedback regulation; and the two gray dashed lines represent the cell membrane and the nuclear membrane. **(D)** Schematic representation of two negative feedback models for regulating Fus3 gene expression. A and B represent, respectively, two different types of proteins that interact with Fus3 in the signaling pathway; “α” stands for α-factor pheromone. **(E)** The fluorescence intensity trajectories of Fus3 inside and outside the nucleus are over time. The red line represents the fitting line of the high state and the low state.

To comprehend the logical relationship between the cell fate at expression levels of Fus3 and pheromone-influenced signaling in the transduction process, we explored the biological mechanism underlying the emergence of these two Fus3 fates. We proposed that the two fates of Fus3 expression observed in signaling could be explained by a logical sequence involving functional depletion and feedback loops. First, functional depletion means that when the amount of Fus3 in the nucleus was sufficient to arrest the cell cycle and activate the mating protein (“Outer_P1 ➔ Far1 & Ste12”), the excess Fus3 was transported out of the nucleus for polar growth (“Outer_P1 ➔ Inner_P1” and “Inner_P2 ➔ Outer_P2”) **(Fig. 3C)**. Second, the feedback loop means that yeast actively regulated Fus3 expression by activating or inhibiting signals. Previously, experimentally validated feedback loops were categorized into two models: “A → Fus3 ⊣ A” focuses primarily on the pathways involving Gpa1 and Sst2, whereas “Fus3 → B ⊣ Fus3” focuses primarily on the pathways involving Ste12 and Msg5 **(Fig. 1 and Fig. 3D)**.

This dynamic sequential mechanism results in two behaviors: the highly coordinated expression levels of Fus3 inside and outside the nucleus, and the alternation of the two fates. As experimental evidence for our proposed molecular mechanism, we observed the existence of two behaviors. We revealed a correlation between intra- and extra-nuclear fluorescence intensities using Pearson’s coefficient. A linear correlation of approximately 0.80 between these two types of fluorescence trajectories at various pheromone concentrations indicated a strong synergy between them **(Table 1)**. In addition, using a hidden Markov chain model, we characterized the two cellular fates using gene expression levels by fitting the fluorescence trajectories into high states (H_1_, H_2_) and low states (L_1_, L_2_) **(Fig. 3E)**. The fitted red line in **Fig. 3E** demonstrates that the two trajectory fates (high and low states) inside and outside the nucleus could switch to each other, supporting our logical interpretation of the state switching above.

**Table 1.**
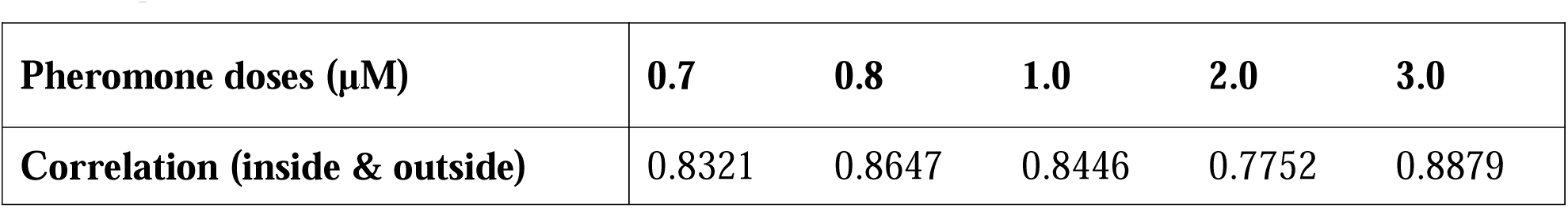
Correlation coefficients of fluorescence trajectories. The “Pearson Correlation Coefficient” between the intranuclear and extranuclear fluorescence intensity trajectories at various pheromone concentrations.

| Pheromone doses ($\mu\text{M}$) | 0.7 | 0.8 | 1.0 | 2.0 | 3.0 |
| --- | --- | --- | --- | --- | --- |
| Correlation (inside & outside) | 0.8321 | 0.8647 | 0.8446 | 0.7752 | 0.8879 |

### Uncovering the physical characteristics of cellular decision-making landscapes

To uncover the physical characteristics of the underlying bistable landscapes, we employed a Hidden Markov Chain model to compute the transition probability, transition rate, and residence time between the two gene expression fates. In the non-equilibrium steady state, the steady state probability can be used to quantify the population landscape P or the potential landscape u (*81, 113–115*), where u is defined as the negative logarithm of the steady state distribution P of gene expression, u= -tn P. As can be seen, the population and potential landscapes (P and u) at various pheromone concentrations have two basins of attractions that are of the two expressions of *FUS3*, high state (H_1_, H_2_) and low state (L_1_, L_2_), can be determined separated by barriers **(Fig. 3B and Fig. S3–S5).** The transition probabilities and residence times through the statistical analysis of the experimental data fitted by the hidden Markov chain model. The transition rate can be derived from the transition probability, whereas the barrier height is determined by fitting two peaks into the landscape. The barrier height is determined by the depth of the basin on the potential energy landscape (u= -tn P) obtained directly from the statistical histogram (P) of the fluorescence signals of the gene expressions **(Table 2)**.

**Table 2.**
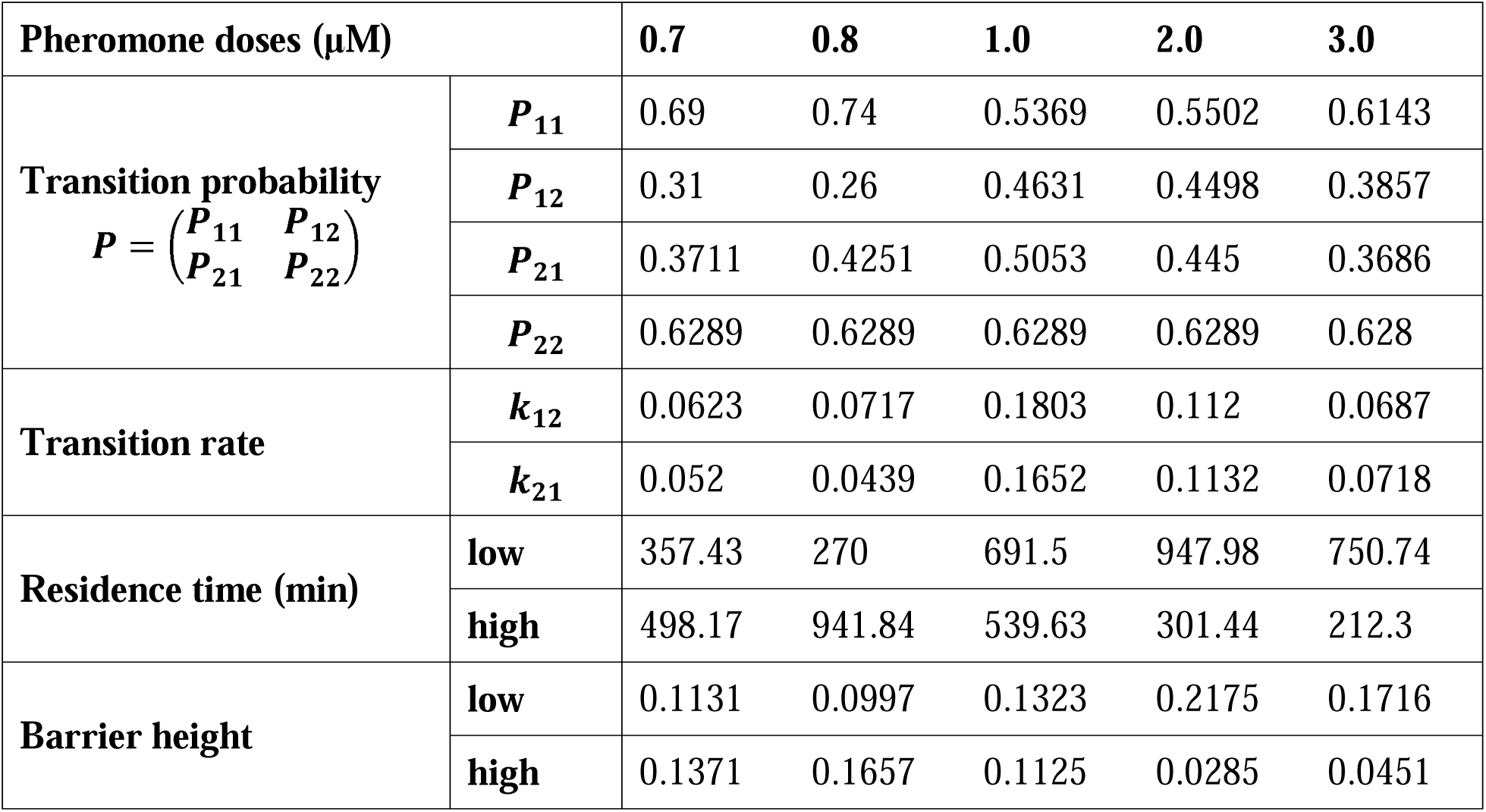
Physical characterization of gene expression landscapes. The transition probability, transition rate, residence time, and barrier height between the high and low states at various pheromone concentrations.

At pheromone concentrations between 0.7 μM and 0.8 μM, the barrier heights of the low state (L_1_, L_2_) were less than those of high state (H_1_, H_2_), whereas the opposite was true between 1.0 μM and 3.0 μM **(Fig. 4A)**. In physics, the transition between steady states becomes harder as the barrier height increases, resulting in a longer residence time (*113, 116*). According to the statistical results of the experimental data, the distribution of high and low states of residence times in different pheromone concentrations is consistent with the distribution of residence times **(Fig. 4B)**. The correlation coefficient between barrier heights and residence times for all concentrations was 0.8806, indicating that they were significantly positively correlated **(Fig. 4C)**. This substantiates the statistical non-equilibrium physics claims regarding barriers and residence time.

**Fig. 4.**
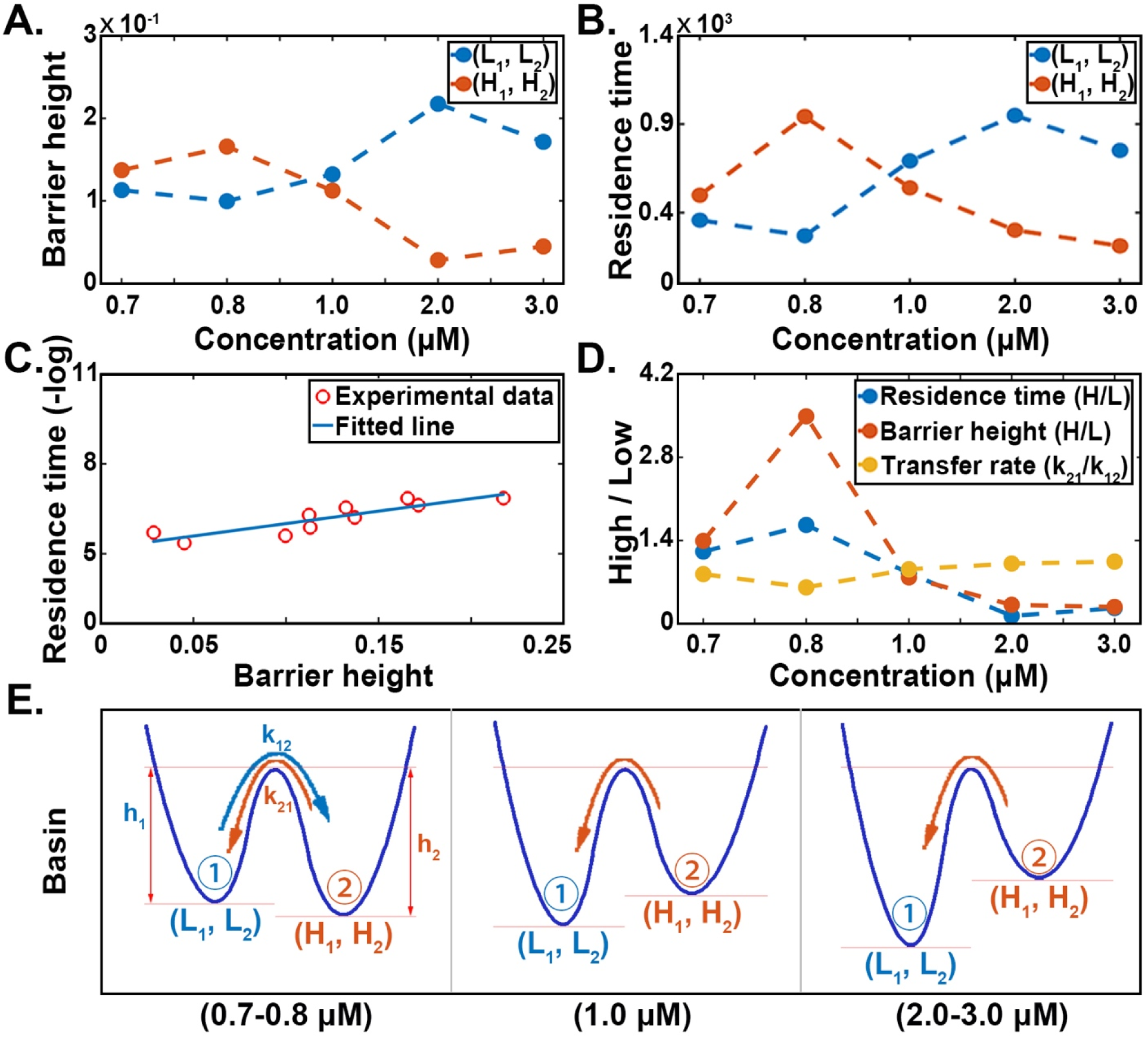
The physical characteristics of cellular decision-making landscapes. **(A)** Trends in the statistical distribution of barrier heights for high and low states at varying pheromone concentrations. **(B)** Trends in the statistical distribution of residence time for high and low states at varying pheromone concentrations. **(C)** Correlation analysis between residence time and barrier height. The red circles are the data points; the blue line is the fitted line of the data. **(D)** The ratio of the physical characteristics of high and low states at various concentrations. The k_12_ is the transition rate from a low state to a high state; k_21_ is the transition rate from a high state to a low state. **(E)** Simple schematic diagram of the potential landscape topography under different pheromone dosages. l1J stands for the low state, l1J stands for the high state; k_12_ and k_21_ are the transition rates between the low state and the high state, respectively; h_1_ and h_2_ represent the respective barrier heights of the low and high states, respectively.

To explore the biological implications of the landscape physical characteristics, we calculated the distribution of their ratios for two gene expression fates as a function of pheromone concentration. The opposite trend of the transfer rate ratio (k_21_/k_12_) relative to the barrier height ratio or the residence time ratio (high/low, H/L) in response to changes in pheromone concentration indicated that transition rates were lower the deeper the attraction basin **(Fig. 4D)**. To better comprehend this relationship, the potential landscape topographies under varying pheromone concentrations were illustrated **(Fig. 4E)**. Compared to a pheromone concentration of 0.7 μM, 0.8 μM had a deeper attraction basin for its high state than for its low state, making the transition from low to high states easier than that from high to low states. The yeast’s cellular decision-making at the level of gene expression was consequently more likely to remain in the high state than in the low state. At 1.0–3.0 μM, the attraction basin of the low state was deeper than that of the high state, leading to the yeast’s preference to remain in the low state. In general, as pheromone doses increase from low to high, the basin of the high state gradually becomes shallower, whereas the basin of the low state gradually becomes deeper. The increased height of the low-state potential barrier is due to the negative feedback regulation that is able to inhibit the expression of Fus3, resulting in a greater number of yeast cells in a low-expression state. This supports the biological notion that high-dose pheromone elicits a stronger negative feedback response compared to the low-dose pheromone.

### Quantifying the cellular deformation rate in polar growth

Upon observing the polar growth of yeast cells in real time via microscopy, we noticed that the rate of cell deformation did not vary uniformly over time. Meanwhile, except for the top, other parts of the cell changed slightly as the yeast stretched forward **(Movies S3–S7)**. To accurately measure the spatiotemporal change rate of the cell morphology, we used a circular filling pattern to segment the yeast cells **(Fig. 5A)**. To quantify the various cell deformations that occur during cell growth, we considered a value (H_n_) comparable to the harmonic mean to characterize the cell morphology. H_n_ equals the sum of the reciprocal radii of the filled circles multiplied by the number of filled circles, i.e., 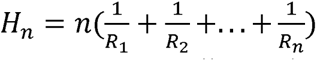. The significant advantage of this parameter is that it is particularly sensitive to small morphological changes at various locations of the cell, allowing differential non-directional polar growth and directional normal growth (i.e., lateral growth and longitudinal growth) to be characterized in real time.

**Fig. 5.**
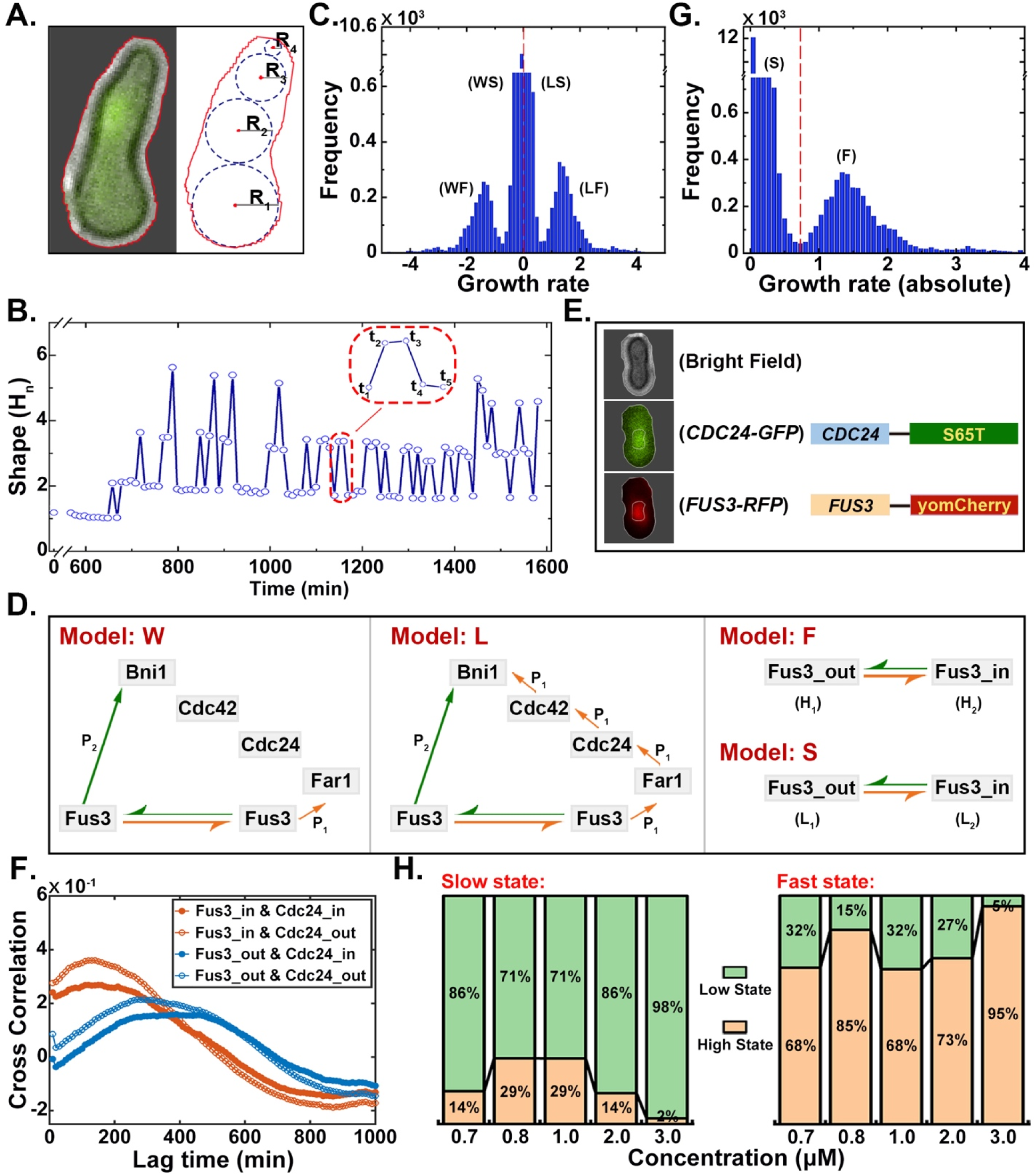
Quantification of the deformation of yeast cells during polar growth. **(A)** A simple diagram of cell shapes with circle filling patterns. The image on the left depicts a yeast cell cultured for 1,020 min in a medium containing 0.7 μM pheromone; the red line indicates the contour of the cell; the image on the right is the filling model for the image on the left; R_1_ - R_4_ are the radii of the circle. **(B)** Real-time trajectory of the cell morphology (H_il_) at 0.7 μM. **(C)** The distribution statistics of the cell growth rate at 0.7 μM. The red dashed line serves as the dividing line between positive and negative data. WF and LS represent the lateral-fast and lateral-slow rates, respectively; LF and WS represent the longitudinal-fast and longitudinal-slow rates, respectively. **(D)** Schematic illustration of the molecular mechanism underlying the formation of the four growth rates. Model W indicates that the polar growth pathway P_1_ is not yet connected, so only the P_2_ pathway operates; Model L depicts the cooperative operation of polar growth pathways P_1_ and P_2_. Models F and S describe the polar growth patterns of the two growth forces, which correspond to the high and low states of Fus3, respectively. **(E)** Dual fluorescent protein system in yeast. The three images on the left, from top to bottom, depict bright field cells, cells excited at 488 nm, and cells excited at 561 nm. The white circle within the cell represents the nucleus boundary; S65T and yomCherry are the fluorescent proteins that were linked to *CDC24* and *FUS3*, respectively. **(F)** Cross-correlation of two levels of gene expression at distinct positions. Gene_in and gene_out represent the gene expression levels inside and outside the nucleus, respectively. **(G)** Statistical graph of the distribution statistics of the absolute value of the cell growth rate at 0.7 μM. **(H)** The proportion of high-state and low-state data present in high and low states at various pheromone concentrations.

From the real-time trajectory of the changes in the cell morphology, as measured by the H_il_ of the cell-filled circles, we found that there were roughly four types of cell shape changes. According to the definition of H_il_ describing the cell morphology, the rising phase (t_1_ - t_2_) of the curve primarily represents the change in cell length, whereas the falling phase (t_3_ - t_4_) primarily represents the change in cell width. At the (t_2_ - t_3_) and (t_4_ - t_S_) phases, the slopes of the two curves are close to zero, indicating that cell morphology changes were minimal or nonexistent **(Fig. 5B and Fig. S14)**. We collected the statistics on the distribution of the cell growth rates or deformation rates **(Fig. 5C and Fig. S6)**. When the positive and negative values were differentiated, it was evident that the cell growth rate could be divided into four states: the lateral-fast rate (WF), the longitudinal-fast rate (LF), the lateral-slow rate (WS) and the longitudinal-slow rate (LS). Among them, L and W represented the presence and absence of direction in cell deformation, or the length and width, respectively. F and S represent the magnitude of the “driving force” in cell deformation, or the fast and slow growth rates, respectively.

To explore the molecular mechanisms underlying these four growth rates, we took both the direction (L and W) and force (F and S) of cell growth into account. Many studies have shown that Fus3 and Far1, which determine the direction of cell growth along its long axis, are essential genes for bud formation during polar cell growth, and their absence results in mis-localization of shmoo projections (*90, 91, 93, 94, 117*). Thus, the indirect pathway (P_1_: “Fus3 → Far1/Cdc24 → Cdc42 → Bni1”), which is more directional than the direct pathway (P_2_: “Fus3 → Bni1”), is the leader signaling pathway for longitudinal cell growth. Given that the P_1_ pathway requires the be transported into or out of the nucleus, we proposed that the P_2_ pathway had a quicker response sequential activation of multiple proteins and that proteins such as Fus3, Far1, and Cdc24 must speed to stimulate cell growth than did the P_1_ pathway (**Fig. 5D**). Consequently, the temporal delay in the actions of these two pathways resulted in P_2_-dominated undirected growth (model W: P_2_), followed by the two growth pathways co-stimulating the cell’s polar growth (model L: P_1_+P_2_) once the P_1_ pathway was fully functional. In addition, we proposed that F and S corresponded to the high and low states in Fus3, respectively. When Fus3 was at a high state (H_1_, H_2_), there was enough Fus3 to stimulate growth (model F). Following entry into the low state (L_1_, L_2_), only a relatively small amount of Fus3 was available for polar growth (model S).

To confirm that there was indeed a time lag in multi-level protein signaling, we constructed a dual fluorescence system (*CDC24*_GFP*-FUS3*_RFP strain), in which *CDC24* was linked to the green fluorescent protein (S65T) and *FUS3* was linked to the red fluorescent protein (yomCherry) **(Fig. 5E and Movies S8–S9)**. Taking into account the order of signal transduction in yeast in response to pheromone, the expression levels of Fus3 and Cdc24 would achieve the highest degree of correlation or match after a certain time lag. Using a cross-correlation function, we analyzed the in- and out-of-nucleus fluorescence trajectories of Fus3 and Cdc24 to compare the lag times of various activation sequences **(Fig. 5F)**. The figure shows that the lag time between Fus3 in the nucleus and Cdc24 (red curve) reaching a maximum correlation was approximately 120 min, while the lag time between Fus3 outside the nucleus and Cdc24 (blue curve) reaching a maximum correlation was approximately 300 min. The fact that the lag time of Fus3_in and Cdc24_in/out (red curve) in the figure was shorter than that of Fus3_out and Cdc24_in/out (blue curve) confirms the existence of a temporal delay effect in multi-level signaling and provides direct evidence for the temporal distinction in the functioning of the two growth pathways.

By taking the growth rate absolute value, we divided the four states into fast growth rate and slow growth rate categories **(Fig. 5G and Fig. S7)**. To confirm that the two forces (F and S) corresponded to the high and low states of Fus3, respectively, the cell growth rate states and Fus3 gene expression states were measured as the cell polar growth changed over time. According to the proportion of statistics in the figure, the High (expression) state was primarily contained within the Fast (growth) state, whereas the Low (expression) state was primarily contained within the Slow (growth) state **(Fig. 5H and Table 3)**. Consequently, this also provided quantitative experimental evidence for the cellular growth rate molecular mechanism.

**Table 3.** The proportional distribution of high state (n_1_, n_2_) and low state (L_1_, L_2_) in two growth rates. “*FH* ” denotes the “High state” in the “Fast growth rate”; “*FL* ” denotes the “Low state” in the “Fast growth rate”; “*SH* ” denotes the “High state” in the “Slow growth rate”; “*SL*” denotes the “Low state” in the “Slow growth rate”; “*FH* + *FL*” and “*SH* + *SL*” represent fast and slow growth rate data volumes.

| **Pheromone doses ($\mu\text{M}$)** | $\frac{FH}{(FH + FL)}$ | $\frac{FL}{(FH + FL)}$ | $(FH + FL)$ | $\frac{SH}{(SH + SL)}$ | $\frac{SL}{(SH + SL)}$ | $(SH + SL)$ |
| --- | --- | --- | --- | --- | --- | --- |
| **0.7** | 0.6847 | 0.3153 | 13888 | 0.1433 | 0.8567 | 7447 |
| **0.8** | 0.8500 | 0.1500 | 5027 | 0.2879 | 0.7121 | 13381 |
| **1.0** | 0.6782 | 0.3218 | 1157 | 0.2872 | 0.7128 | 21884 |
| **2.0** | 0.7266 | 0.2734 | 2249 | 0.1353 | 0.8647 | 34637 |
| **3.0** | 0.9530 | 0.0470 | 1679 | 0.0244 | 0.9756 | 16597 |

### Quantifying the four morphological fates and the phase transition trend

The morphological trajectory of the cell shape over time as described by H_n_ fluctuated continuously within a given range **(Fig. 5B)**. Using a hidden Markov chain model to fit the time-varying trajectories of cell morphology, yeast cells at different pheromone concentrations exhibited four distinct morphological fates. These four morphological fates corresponded statistically to the four peaks in the distribution of all cell shapes (F_1_–F_4_) **(Fig. 6A and Fig. S8)**. Due to the fact that H_n_ is a parameter that is extremely sensitive to changes in the size and number of filled circles within the cell, minute variations in the length or width of a portion of the cell observed through a microscope can result in the cell being classified as having a different morphological fate **(Fig. 6B)**. We proposed, on the basis that cell morphology is an accumulation in growth, that the molecular mechanisms underlying the formation of the four morphological fates were dependent on the different capacities of cells to grow laterally and longitudinally. To test the claim that four morphological fates were formed, the synergistic effect of growth ability in both directions was calculated. The lateral growth capacity of cells was expressed by the rate of change of the average radius of the filled circle within the cell, whereas the longitudinal growth capacity was expressed by the rate of change of the sum of the radii of the filled circle. The existence of four distinct states in the cooperative distribution of the two data, as depicted in the figure, strongly suggested that the capacity to grow in different directions is a crucial factor in determining cell morphology **(Fig. 6C and Fig. S9)**.

**Fig. 6.**
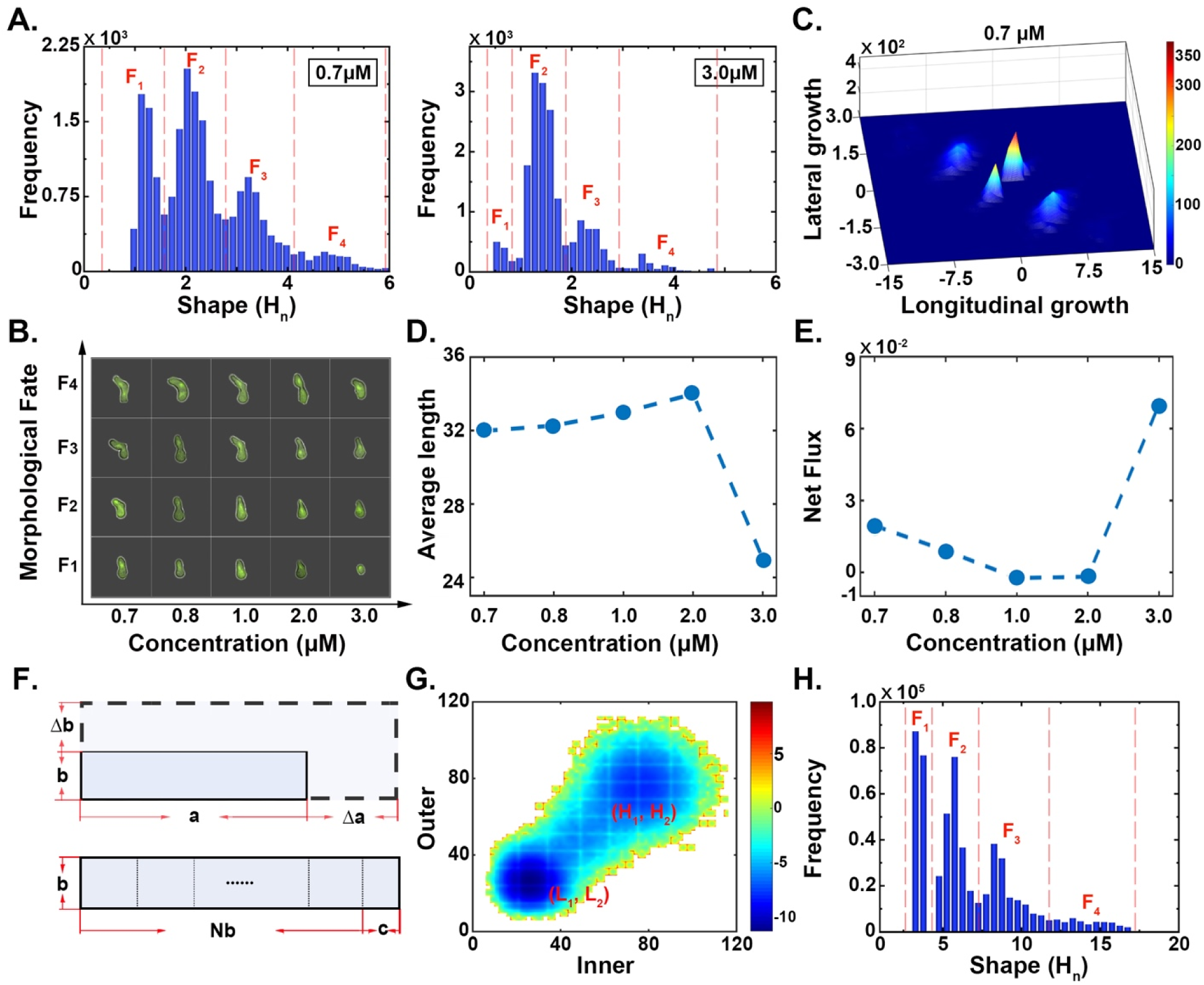
The interpretation of the different cell morphological fates. **(A)** Statistical distribution of the cell morphology at 0.7 μM. The red dashed lines roughly correspond to the boundary between distinct cell morphological fates; F_1_–F_4_ represent the four cell morphological fates. **(B)** Photographs taken with a fluorescence microscope of cells exhibiting four distinct morphological fates in response to varying pheromone concentrations. **(C)** The synergistic effect of both lateral and longitudinal cell growth capabilities at a pheromone concentration of 0.7 μM. The x-axis represents the ability of the cells to grow longitudinally, as indicated by the rate of change in the sum of the radii of the filled circles within the cells, i.e., cR_1_ + R_2_ +·+ R_il_)′; the y-axis represents the ability of cells to grow laterally, as indicated by the rate of change in the average radius of the filled circles within the cells, i.e., 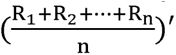. **(D)** Changes in the average cell _il_ length as a function of pheromone concentration. **(E)** The sum of the net fluxes among the four growth; a represents the length of the cell; Δa is the increased length of the cell; b represents the cell morphologies at various pheromone concentrations. **(F)** The scheme for the simulated cell width of the cell; and Δb is the increased width of the cell; the scheme of the H_n_ calculation; c is the remainder of the cell length divided by b; N represents the number of b. **(G)** Distribution graph of the negative log of the value of the Fus3 fluorescence intensity inside and outside the nucleus of yeast cells in the stationary phase using simulation. **(H)** The distribution of the cell morphology (H_n_) using simulation.

Notably, a phase transition trend from four states to a dominant state in the cell morphology distribution between 0.7 μM and 3.0 μM was revealed (**Fig. 6A**). Moreover, the morphology of the cells observed under a microscope at a concentration of 3.0 μM was significantly smaller than that observed at other concentrations **(Fig. 6B)**. By measuring the average cell length, it is possible to conclude that a high-dose pheromone concentration (3.0 μM) disrupted the monotonic linear trend observed at low doses (0.7 μM-2.0 μM) **(Fig. 6D and Table 4)**. The microscopic explanation of this phase transition trend is the enhancement of negative feedback regulation under high doses (I_2_), suppressing the second polar growth pathway (P_2_) **(Fig. 1 and Fig. 3C)**. Three-quarters of the cell fates (with the exception of F_2_) showed a sudden reduction in morphology at a high pheromone dose (3.0 μM), most likely attributed to this impaired P_2_ function **(Fig. 6A and Fig. S8)**.

**Table 4.** The Average length of yeast cells at different pheromone doses.

| **Pheromone doses ($\mu\text{M}$)** | **0.7** | **0.8** | **1.0** | **2.0** | **3.0** |
| --- | --- | --- | --- | --- | --- |
| **Average length** | 31.98 | 32.27 | 32.99 | 33.99 | 25.01 |

To explain the physical mechanism of the phase transition trend that occurs at high dose (3.0 μM), we quantified the degree of nonequilibrium in the cellular morphological system using the net flux. From the cell morphological trajectories analyzed by the hidden Markov chain model, we determined the transition probability between various cell morphologies. The sum of the three net fluxes, which was obtained by decomposing the probability loops in the transition matrix, was used to represent the degree of the detailed balance collapsed in the cellular morphological system **(Fig. 6E)**. As the concentration of pheromone increased, the intensity of the net flux decreased first and then increased, exhibiting a significant phase transition trend at 3.0 μM. Since the net flux is rotational due to its steady state nature under local probability conservation, it tends to destabilize the point attractors. Therefore, the significant changes in net flux can lead to the instability of the cell attractor states, giving rise to possible phase transition trend. The quantification of this non-equilibrium dynamic explains the physical mechanism by which the phase transition trend in a morphological system was caused by the enhanced capability of the yeast cells negative feedback regulation at high doses.

### Simulations for the signal transduction and the cell growth

Based on the functional and quantitative regulation obtained from databases, such as KEGG (Kyoto Encyclopedia of Genes and Genomes, https://www.kegg.jp/), SGD (the *Saccharomyces* Genome Database, https://www.yeastgenome.org/) and EVEX (http://evexdb.org/), we developed a simplified model of signal transduction in the context of global gene regulatory networks **(Fig. 1, Fig. 6F)**. This model simulated a series of biochemical reactions with the Gillespie algorithm (*118–120*) **(Table S6–8)**. By simulating the biochemical reactions associated with the pheromone pathway in yeast cells, the expression levels of Fus3 and the distribution of cell morphology were determined. Fus3 gene expression obtained from biochemical reaction results displayed a two-state distribution inside and outside of the nucleus, validating our understanding of the molecular mechanism underlying the bimodal fluorescence state within the context of the global response **(Fig. 6G)**. Meanwhile, the Bni1 produced by the reactions simulated the dynamic process of cell growth. Although both the growth pathways (P_1_ and P_2_) were involved in the process of cellular length and width growth, the relative weights of the pathways that grew in the two directions were significantly different. Therefore, we simply defined Bni1_in produced by the P_1_ pathway that plays a major role as the longitudinal growth and Bni1_out produced by the P_2_ pathway as the lateral growth.

In this simulation, a denotes the longitudinal length of the yeast cell, while b denotes the lateral length of the yeast cell **(Fig. 6F)**. Each iteration increased the longitudinal and lateral lengths of the yeast cells by Δa and Δb, respectively. Δa is proportional to “PBni1_in” and inversely proportional to b; Δb is proportional to “PBni1_out” and inversely proportional to a. That is, 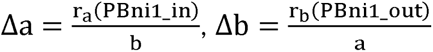. The value N corresponds to the rounding operation of 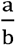, i.e., 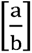, and c represents the amount remaining after rounding. The H_n_ is cN + 1)cN⁄b + 1⁄c). As predicted by the simulation of the cell growth process, the distribution of cell morphology revealed four distinct fates, which corresponded to the experimental observations **(Fig. 6H)**.

## DISCUSSION

In this study, we quantitatively uncovered and interpreted the yeast cell fate decision-making in response to pheromone using biological and physical methods. Using flow cytometry, we examined the induction of yeast cells by different concentrations of pheromone to determine the critical threshold for eliciting a cellular response. The purpose of the fitting curve in **Fig. 2A** was to estimate the pheromone dose by determining at what concentration the expression of Fus3 exhibited double peaks. Continuous microscopy observations of *FUS3*-GFP strain yeast cells provided us with a microscopic and mesoscopic view of how the cells responded to pheromone. The four stages of Fus3 expression levels at a 0.7 μM pheromone concentration accurately depicted the mesoscopic cell behavior of cell cycle arrest and polar growth (**Fig. 2B**). Among them, the Fus3 expression levels in the stable stage (b2–b3) fluctuate around a fixed value, which was considered a non-equilibrium steady state period. As our subsequent analysis centered on the non-equilibrium steady-state phase, we chose five pheromone concentrations as stimuli that allowed the yeast cells to maintain a stable-state phase for a sufficient amount of time.

To explore the fate of Fus3 gene expression in a non-equilibrium steady state, we chose data that, after 600 min, brought all the cell fluorescence trajectories into stable-state phase **(Fig. 3A)**. The two fates of Fus3 that resulted from cellular decision-making were separated by the Markov fitting of the trajectories **(Fig. 3B)**. The criterion for data fitting is the probability that the fluorescence intensity at a particular moment in the trajectory belonged to a high state (H_1_, H_2_) or a low state (L_1_, L_2_) multiplied by their respective transition probabilities **(Fig. 3E)**. Consequently, there was a chance that the value of the fluorescence intensity in the low state was greater than the value of the fluorescence intensity in the high state. We proposed that the two fates of Fus3 expressions observed in signaling were explained by a logical sequence of point-to-point feedback regulation between proteins. Notably, even though the protein-to-protein feedback regulation described above had been validated by previous biological experiments (*85-89, 95-104*), our claims focused primarily on establishing a dynamical logical connection between these isolated regulatory processes from a global perspective. For the purpose of validating this dynamic molecular mechanism, one may wonder why the (L_1_, H_2_) and (H_1_, L_2_) states did not appear in this scheme. We know from experimental evidence that the absence of the (L_1_, H_2_) and (H_1_, L_2_) states was due to the strong correlation between intra- and extra-nuclear fluorescence intensities. If the four fates emerge on the landscape of intra- and extra-nuclear fluorescence intensities of Fus3, the Pearson correlation coefficient between these two types of trajectories would be zero (*121*). Moreover, the feedback model of forward activation and negative inhibition favored the formation of two states from a physical standpoint **(Fig. 3D)**.

Between steady states, the cellular decision-making is primarily reflected by the transition rate, transition probability, residence time, and potential barrier height. Due to the relative nature of the high and low states at each pheromone concentration, their physical characteristics can be compared directly, but not at different concentrations. To compare the relative significance of these physical characteristics at different concentrations, a similar normalized approach (high/low) was used. At 0.7–0.8 μM, a slight increase in barrier height significantly lengthened the residence time of the high state. At 1.0–3.0 μM, the residence time also decreased gradually as the barrier height decreased **(Fig. 4D)**. In our biological system, the positive correlation statement regarding barrier height and residence time was confirmed **(Fig. 4C)**. Certainly, such physical characteristics that quantify cellular decision-making have significant biological implications for the study of yeast response behavior. Positive feedback regulation in the gene network (“Pheromone → Fus3” and “Fus3 → Fus3”), for example, increased the potential barrier height on the gene expression landscape, whereas negative feedback regulation decreased it (I_1_, I_2_) **(Fig. 1 and Fig. 3C)**. The residence time at various pheromone concentrations revealed which gene expression fate yeast cells preferred in response to the pheromone.

Some studies believe that the establishment of polarity helps organisms to survive better in nature (*122, 123*). Due to the inability of yeast’s chemotactic response to swim as that of E. coli, its polar growth could only grow in the direction of a high concentration of pheromone, achieving mutual contact and fusion between heterosexual yeasts (*4, 124, 125*). To quantify the mesoscopic behavior of yeast cells in polar growth, which grew and stopped intermittently, the yeast cells were characterized as a combination of filled circles. The white edge of the yeast cell image was the cell wall **(Fig. 5A).** Therefore, the cell filling circles were based on the outermost edge of the cell wall as the cell boundary. Rather than using the cell area, the total length, or the arithmetic mean of the length, we chose to describe the cell morphology using the reciprocal of the radius in the cell-filled circle multiplied by the number of filled circles, i.e., 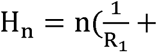 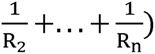. This method has the following advantages: first, it could reflect the deformation of different parts of the cell; and second, H_n_ is a statistic that can reflect both the process and outcome of the cell polarity growth. It was important to note that when a cell grew laterally, its length also increased, so the terms lateral growth and longitudinal growth were relative rather than absolute descriptions of length and width.

The cell growth rate primarily characterized the process of cell polar growth, whereas the cell morphology primarily characterized the outcome. Specifically, the cell growth rate is actually the cell deformation rate. To verify the molecular mechanism of the growth rate for directional (model L) and non-directional (model W) growth, we constructed a dual fluorescence system (*CDC24*_GFP*-FUS3*_RFP) to demonstrate that there was a time lag in the multilevel protein signaling behavior. Due to the different maturation times of the two fluorescent proteins (GFP and RFP), the lag time between Fus3 and Cdc24 was only a relative quantity. Nonetheless, the delay time of the dual-fluorescence system at different positions was used to quantify the practical significance of multipolar signal transduction. Notably, the non-directional growth (model W: P_2_) did not consist solely of the polar growth pathway of P_2_, but rather a relative weight simplification to differentiate it from the directional growth (model L: P_1_+P_2_). To confirm that the fast growth rate (model F) and slow growth rate (model S) corresponded to the high and expression states. The high state (H_1_, H_2_) was primarily included in the fast growth rate, whereas low states of Fus3, we explored the association between growth rates and dual fluorescent gene the low state (L_1_, L_2_) was primarily included in the slow growth rate, confirming the molecular mechanism of F and S **(Fig. 5H and Table 3)**. In addition, there must be a lag between the expression of Fus3p and the observed cell growth, resulting in a low fluorescence state during fast cell growth and a high fluorescence state during slow cell growth.

For statistical analysis of different cell morphologies, we distinguished four cell morphological fates roughly as F_1_-F_4_ to provide a more intuitive description **(Fig. 6A)**. Nevertheless, the division of actual cell morphological fate was determined by fitting the trajectories, and not arbitrarily by the size of H_n_. Given that H_n_ is extremely sensitive to changes in length and width in different parts of the cell, the area of cells that were morphologically divided into the four categories did not differ significantly from their overall appearance **(Fig. 6B)**. Our justification for proposing the molecular mechanisms of the four cell morphological fates is that morphology is the accumulation of a growth process. Due to the fact that the cell morphology described by H_il_ resembled the relative length and width of a cell, the capacity to grow in both directions can explain the emergence of cell morphologies **(Fig. 6C)**. In addition, in the statistical graph of cell morphology, other fates tended to disappear, with the exception of the increase in the proportion of F_2_. This indicated that the intracellular phase transition trend occurred at 3.0 μM as a result of enhanced negative feedback regulation (I_2_ in **Fig. 1**). The coordinates of the cell morphology statistics chart at 3.0 μM were deviated from those of other concentrations due to the phase transition trend in the system, which altered the measurement scale of the four morphological fates. To explain the physics of the phase transitions in the biological system, we quantified the degree to which the detailed balance collapsed by employing the net flux characterizing the non-equilibrium statistical physics. The results demonstrated that the non-equilibrium dynamics of the biological system was supported by the significant increase in net flux of the morphological system at high pheromone dose (3.0 μM), causing the instability and phase transition.

Finally, we developed a simplified model of the gene regulatory network for signal transduction in order to confirm the global rationality of the dynamic sequential mechanism. Thus far, we have established the logical links between the functions of regulated proteins in the pheromone pathway for mating through the underlying signal transduction process. While our validation method differs from the conventional approach of constructing yeast mutants, it has been successful in observing realistic cellular response behavior. Furthermore, our future research will focus on utilizing fluorescent labels to enable real-time visualization of the dynamics of signaling pathways. Specifically, we plan to construct a multi-dimensional fluorescent system to elucidate the temporal and spatial dynamics of cellular responses, including potential crosstalk between different signaling pathways. This approach will provide a more comprehensive and quantitative understanding of how cells respond to various stimuli, and how these responses are integrated at the molecular level.

## MATERIALS AND METHODS

### Yeast strains and Growth media

The yeast strain used in this experiment is *Saccharomyces cerevisiae* S288C (ATCC 201388: *MAT*a*his*3Δ1 *leu*2Δ0 *met*15Δ0 *ura*3Δ0) (*126*). In the yeast fluorescence library, we choose a special yeast as the research object, whose c-terminal of Fus3 is fused with GFP as the reporter protein. The yeasts are cultured by inoculating 5ml of YNB [Yeast Nitrogen Base Without Amino Acids (6.8 g/L), Dextrose (5 g/L), Uracil (76mg/L), 50×MEM Amino Acids (20ml/L)] medium with a colony from a YPD agar plate [Yeast extract (10 g/L), Peptone (20 g/L), Dextrose (20 g/L), Agar (10 g/L)]. Cells are grown overnight at 30LJ shaking at 250 rpm (about 14h). Before starting the experiment, we take out 20ul of overnight cultures and dilute 100-fold in 2ml YNB medium to obtain an optical density of 0.1 at 600 nm (OD600nm).

### Flow cytometry

To determine the critical concentration of pheromone, we chose a flow cytometer for preliminary screening. We set 8 different concentrations (0.01μM, 0.05μM, 0.1μM, 0.2 μM, 0.4μM, 0.6μM, 1.0μM, 1.5μM) for the pheromone medium, and culture the yeast cells in a shaker at 250 rpm and 30LJ. In order to avoid contamination of the yeast cells, we have set up 5 sets of parallel culture samples for each concentration of the medium. After culturing on a shaker for specific times (3h, 6h, 9h, 12h and 24h), 105,000 cells are taken from each sample for analysis using the flow cytometer.

### Microfluidic system platform

We use the 4-chamber cell culture plate, CellASIC™ ONIX Y04C-02 Microfluidic Plate, for live cell culture at 30°C. The plate is designed for use with the CellASIC™ ONIX Microfluidic System, which provides a controlled dynamical microenvironment for cells. First, the yeast cells are cultured in a microfluidic plate containing conventional YNB medium for 1 hour to adapt to the new environment, and then switch to YNB medium supplemented with pheromone and culture at a flow rate of 1 psi for 30 hours.

### Microscopy Measurements

The fluorescence values of the single cells were measured using an inverted fluorescence microscopy (Ti-E, Nikon) with automated stage and focus, equipped with a high NA oil-immersion objective (1.45NA, 100×). We applied 488nm laser and set the output power at 30mW (only 10% of the laser beam into the microscope objective), the fluorescence signals were collected by a cooled EM-CCD camera (897U, Andor). All images were acquired using both bright field imaging and fluorescent field imaging. These images were acquired by Nikon software. Data analysis were accomplished through a combination of manual and automated analysis using custom Matlab code. Many trajectories were taken from a time-lapse microscopy. The fluorescent images were periodically captured and recorded every 10 minutes. The fluorescent data of each cell at each time point were collected for the following discussion.

## Acknowledgments

SL, QL, and EW acknowledge the support of the National Natural Science Foundation of China with Grant No. 21721003.

## Data and materials availability

All data are available in the main text or the supplementary materials.

## References and Notes

1. H. C. Berg, D. A. Brown, Chemotaxis in Escherichia coli analysed by three-dimensional tracking. Nature 239, 500–504 (1972).

2. R. M. Macnab, D. E. Koshland Jr, The gradient-sensing mechanism in bacterial chemotaxis. Proceedings of the National Academy of Sciences 69, 2509–2512 (1972).

3. J. Yuan, K. A. Fahrner, L. Turner, H. C. Berg, Asymmetry in the clockwise and counterclockwise rotation of the bacterial flagellar motor. Proceedings of the National Academy of Sciences 107, 12846–12849 (2010).

4. V. Sourjik, N. S. Wingreen, Responding to chemical gradients: bacterial chemotaxis. Current opinion in cell biology 24, 262–268 (2012).

5. S. De Oliveira, E. E. Rosowski, A. Huttenlocher, Neutrophil migration in infection and wound repair: going forward in reverse. Nature Reviews Immunology 16, 378–391 (2016).

6. P. Zhou et al., Stochasticity triggers activation of the S-phase checkpoint pathway in budding yeast. Physical Review X 11, 011004 (2021).

7. B. Fu et al., Metal-induced sensor mobilization turns on affinity to activate regulator for metal detoxification in live bacteria. Proceedings of the National Academy of Sciences 117, 13248–13255 (2020).

8. J. E. Slessareva, H. G. Dohlman, G protein signaling in yeast: new components, new connections, new compartments. Science 314, 1412–1413 (2006).

9. R. E. Chen, J. Thorner, Function and regulation in MAPK signaling pathways: Lessons learned from the yeast Saccharomyces cerevisiae. Bba-Mol Cell Res 1773, 1311–1340 (2007).

10. Y. Q. Wang, H. G. Dohlman, Pheromone signaling mechanisms in yeast: A prototypical sex machine. Science 306, 1508–1509 (2004).

11. M. C. Gustin, J. Albertyn, M. Alexander, K. Davenport, MAP Kinase Pathways in the YeastSaccharomyces cerevisiae. Microbiology and Molecular biology reviews 62, 1264–1300 (1998).

12. I. Herskowitz, Life cycle of the budding yeast Saccharomyces cerevisiae. Microbiological reviews 52, 536 (1988).

13. C. M. Hull, R. M. Raisner, A. D. Johnson, Evidence for mating of the" asexual" yeast Candida albicans in a mammalian host. Science 289, 307–310 (2000).

14. J. Warringer et al., Trait variation in yeast is defined by population history. PLoS Genet 7, e1002111 (2011).

15. E. Zörgö et al., Ancient evolutionary trade-offs between yeast ploidy states. PLoS Genet 9, e1003388 (2013).

16. D. Norris, M. Osley, The two gene pairs encoding H2A and H2B play different roles in the Saccharomyces cerevisiae life cycle. Molecular and Cellular Biology 7, 3473–3481 (1987).

17. J. E. Haber, Mating-type genes and MAT switching in Saccharomyces cerevisiae. Genetics 191, 33–64 (2012).

18. M. Z. Bao, M. A. Schwartz, G. T. Cantin, J. R. Yates III, H. D. Madhani, Pheromone-dependent destruction of the Tec1 transcription factor is required for MAP kinase signaling specificity in yeast. Cell 119, 991–1000 (2004).

19. D. D. Jenness, A. C. Burkholder, L. H. Hartwell, Binding of α-factor pheromone to yeast a cells: chemical and genetic evidence for an α-factor receptor. Cell 35, 521–529 (1983).

20. H. Youk, W. A. Lim, Secreting and sensing the same molecule allows cells to achieve versatile social behaviors. Science 343, 1242782 (2014).

21. H. Youk, W. A. Lim, Secreting and Sensing the Same Molecule Allows Cells to Achieve Versatile Social Behaviors. Science 343, 628-+ (2014).

22. M. D. Rose, Nuclear fusion in the yeast Saccharomyces cerevisiae. Annu Rev Cell Dev Bi 12, 663–695 (1996).

23. L. Marsh, M. Rose, The pathway of cell and nuclear fusion during mating in S. cerevisiae. COLD SPRING HARBOR MONOGRAPH SERIES 21, 827–888 (1997).

24. J. M. White, M. D. Rose, Yeast mating: getting close to membrane merger. Current biology 11, R16–R20 (2001).

25. L. Bardwell, A walk-through of the yeast mating pheromone response pathway (vol 25, pg 1465, 2004). Peptides 26, 337-+ (2005).

26. J. N. Molk, K. Bloom, Microtubule dynamics in the budding yeast mating pathway. Journal of cell science 119, 3485–3490 (2006).

27. H. G. Dohlman, J. W. Thorner, Regulation of G protein-initiated signal transduction in yeast: Paradigms and principles. Annu Rev Biochem 70, 703–754 (2001).

28. A. M. Neiman, I. Herskowitz, Reconstitution of a Yeast Protein-Kinase Cascade in-Vitro - Activation of the Yeast Mek Homolog Ste7 by Ste11. P Natl Acad Sci USA 91, 3398–3402 (1994).

29. B. Errede, A. Gartner, Z. Zhou, K. Nasmyth, G. Ammerer, MAP kinase-related FUS3 from S. cerevisiae is activated by STE7 in vitro. Nature 362, 261–264 (1993).

30. P. M. P. Anne-Christine Butty, Linda S. Huang, Ira Herskowitz, Matthias Peter*, The Role of Far1p in Linking the Heterotrimeric G Protein to Polarity Establishment Proteins During Yeast Mating. SCIENCE, (1998).

31. D. Matheos, M. Metodiev, E. Muller, D. Stone, M. D. Rose, Pheromone-induced polarization is dependent on the Fus3p MAPK acting through the formin Bni1p. The Journal of cell biology 165, 99–109 (2004).

32. M. Peter, A. Gartner, J. Horecka, G. Ammerer, I. Herskowitz, FAR1 links the signal transduction pathway to the cell cycle machinery in yeast. Cell 73, 747–760 (1993).

33. E. Elion, B. Satterberg, J. Kranz, FUS3 phosphorylates multiple components of the mating signal transduction cascade: evidence for STE12 and FAR1. Molecular biology of the cell 4, 495–510 (1993).

34. R. L. Roberts, G. R. Fink, Elements of a single MAP kinase cascade in Saccharomyces cerevisiae mediate two developmental programs in the same cell type: mating and invasive growth. Genes & development 8, 2974–2985 (1994).

35. I. Herskowitz, MAP kinase pathways in yeast: for mating and more. Cell 80, 187–197 (1995).

36. E. A. Elion, J. A. Brill, G. R. Fink, FUS3 represses CLN1 and CLN2 and in concert with KSS1 promotes signal transduction. Proceedings of the National Academy of Sciences 88, 9392–9396 (1991).

37. E. Blackwell et al., Effect of the pheromone-responsive Gα and phosphatase proteins of Saccharomyces cerevisiae on the subcellular localization of the Fus3 mitogen-activated protein kinase. Molecular and cellular biology 23, 1135–1150 (2003).

38. F. van Drogen, V. M. Stucke, G. Jorritsma, M. Peter, MAP kinase dynamics in response to pheromones in budding yeast. Nature cell biology 3, 1051–1059 (2001).

39. F. Chang, I. Herskowitz, Identification of a gene necessary for cell cycle arrest by a negative growth factor of yeast: FAR1 is an inhibitor of a G1 cyclin, CLN2. Cell 63, 999-1011 (1990).

40. Y. Shimada, M.-P. Gulli, M. Peter, Nuclear sequestration of the exchange factor Cdc24 by Far1 regulates cell polarity during yeast mating. Nature cell biology 2, 117–124 (2000).

41. J. E. Segall, Polarization of yeast cells in spatial gradients of alpha mating factor. Proceedings of the National Academy of Sciences 90, 8332–8336 (1993).

42. X. N. Fang et al., Cell fate potentials and switching kinetics uncovered in a classic bistable genetic switch. Nature Communications 9, (2018).

43. Z. L. Jiang et al., The emergence of the two cell fates and their associated switching for a negative auto-regulating gene. Bmc Biology 17, (2019).

44. P. A. Takizawa, A. Sil, J. R. Swedlow, I. Herskowitz, R. D. Vale, Actin-dependent localization of an RNA encoding a cell-fate determinant in yeast. Nature 389, 90–93 (1997).

45. A. Colman-Lerner et al., Regulated cell-to-cell variation in a cell-fate decision system. Nature 437, 699–706 (2005).

46. R. M. Gordley et al., Engineering dynamical control of cell fate switching using synthetic phospho-regulons. P Natl Acad Sci USA 113, 13528–13533 (2016).

47. Z. Jiang et al., The emergence of the two cell fates and their associated switching for a negative auto-regulating gene. BMC biology 17, 1–14 (2019).

48. L. Zeng et al., Decision making at a subcellular level determines the outcome of bacteriophage infection. Cell 141, 682–691 (2010).

49. A. Colman-Lerner et al., Regulated cell-to-cell variation in a cell-fate decision system. Nature 437, 699–706 (2005).

50. X. Fang et al., Cell fate potentials and switching kinetics uncovered in a classic bistable genetic switch. Nature communications 9, 1–9 (2018).

51. F. St-Pierre, D. Endy, Determination of cell fate selection during phage lambda infection. Proceedings of the National Academy of Sciences 105, 20705–20710 (2008).

52. P. J. Choi, L. Cai, K. Frieda, X. S. Xie, A stochastic single-molecule event triggers phenotype switching of a bacterial cell. Science 322, 442–446 (2008).

53. L. S. Weinberger, J. C. Burnett, J. E. Toettcher, A. P. Arkin, D. V. Schaffer, Stochastic gene expression in a lentiviral positive-feedback loop: HIV-1 Tat fluctuations drive phenotypic diversity. Cell 122, 169–182 (2005).

54. W. Wang et al., Live-cell imaging and analysis reveal cell phenotypic transition dynamics inherently missing in snapshot data. Sci Adv 6, eaba9319 (2020).

55. N. Yosef, A. Regev, Impulse Control: Temporal Dynamics in Gene Transcription. Cell 144, 886–896 (2011).

56. Z. Bar-Joseph, A. Gitter, I. Simon, Studying and modelling dynamic biological processes using time-series gene expression data. Nature Reviews Genetics 13, 552–564 (2012).

57. M. Behar, N. Hao, H. G. Dohlman, T. C. Elston, Dose-to-Duration Encoding and Signaling beyond Saturation in Intracellular Signaling Networks. Plos Computational Biology 4, (2008).

58. A. Doncic et al., Compartmentalization of a Bistable Switch Enables Memory to Cross a Feedback-Driven Transition. Cell 160, 1182–1195 (2015).

59. J. Hasty, D. McMillen, F. Isaacs, J. J. Collins, Computational studies of gene regulatory networks: in numero molecular biology. Nature Reviews Genetics 2, 268–279 (2001).

60. F. Caudron, Y. Barral, A super-assembly of Whi3 encodes memory of deceptive encounters by single cells during yeast courtship. Cell 155, 1244–1257 (2013).

61. J. S. Weinberg, D. G. Drubin, Regulation of clathrin-mediated endocytosis by dynamic ubiquitination and deubiquitination. Curr Biol 24, 951–959 (2014).

62. M. D. Leonetti, S. Sekine, D. Kamiyama, J. S. Weissman, B. Huang, A scalable strategy for high-throughput GFP tagging of endogenous human proteins. Proc Natl Acad Sci U S A 113, E3501–3508 (2016).

63. R. G.-L. Patrick Conlon, Ambhighainath Ganesan, Jin Zhang, and Andre Levchenko, Single-cell dynamics and variability of MAPK activity in a yeast differentiation pathway. PNAS, (2016).

64. S. Ozsezen et al., Inference of the High-Level Interaction Topology between the Metabolic and Cell-Cycle Oscillators from Single-Cell Dynamics. Cell Syst 9, 354–365 e356 (2019).

65. A. K. Chakravarty, T. Smejkal, A. K. Itakura, D. M. Garcia, D. F. Jarosz, A Non-amyloid Prion Particle that Activates a Heritable Gene Expression Program. Mol Cell 77, 251–265 e259 (2020).

66. C. L. Klaips, M. H. M. Gropp, M. S. Hipp, F. U. Hartl, Sis1 potentiates the stress response to protein aggregation and elevated temperature. Nature Communications 11, (2020).

67. S. N. Mouton et al., A physicochemical perspective of aging from single-cell analysis of pH, macromolecular and organellar crowding in yeast. Elife 9, (2020).

68. A. K. Pfitzner et al., An ESCRT-III Polymerization Sequence Drives Membrane Deformation and Fission. Cell 182, 1140–1155 e1118 (2020).

69. C. Ptak et al., Phosphorylation-dependent mitotic SUMOylation drives nuclear envelope-chromatin interactions. J Cell Biol 220, (2021).

70. W. Feng, O. Argüello-Miranda, S. Qian, F. Wang, Cdc14 spatiotemporally dephosphorylates Atg13 to activate autophagy during meiotic divisions. Journal of Cell Biology 221, (2022).

71. S. Pinheiro, S. Pandey, S. Pelet, Cellular heterogeneity: yeast-side story. Fungal Biology Reviews 39, 34–45 (2022).

72. S. Chaturvedi et al., Disrupting autorepression circuitry generates "open-loop lethality" to yield escape-resistant antiviral agents. Cell 185, 2086–2102 e2022 (2022).

73. W. Wang, G. W. Li, C. Chen, X. S. Xie, X. Zhuang, Chromosome organization by a nucleoid-associated protein in live bacteria. Science 333, 1445–1449 (2011).

74. H. Feng, J. Wang, Potential and flux decomposition for dynamical systems and non-equilibrium thermodynamics: Curvature, gauge field, and generalized fluctuation-dissipation theorem. The Journal of chemical physics 135, 234511 (2011).

75. C. Li, J. Wang, Landscape and flux reveal a new global view and physical quantification of mammalian cell cycle. Proceedings of the National Academy of Sciences 111, 14130–14135 (2014).

76. G. Balázsi, A. van Oudenaarden, J. J. Collins, Cellular decision making and biological noise: from microbes to mammals. Cell 144, 910–925 (2011).

77. J. Wang, K. Zhang, L. Xu, E. Wang, Quantifying the Waddington landscape and biological paths for development and differentiation. Proceedings of the National Academy of Sciences 108, 8257–8262 (2011).

78. J. Wang, K. Zhang, E. Wang, Kinetic paths, time scale, and underlying landscapes: A path integral framework to study global natures of nonequilibrium systems and networks. The Journal of chemical physics 133, 09B613 (2010).

79. J. Wang, C. Li, E. Wang, Potential and flux landscapes quantify the stability and robustness of budding yeast cell cycle network. Proceedings of the National Academy of Sciences 107, 8195–8200 (2010).

80. J. Wang, K. Zhang, E. Wang, Robustness and dissipation of mitogen-activated protein kinases signal transduction network: Underlying funneled landscape against stochastic fluctuations. The Journal of chemical physics 129, 10B602 (2008).

81. J. Wang, L. Xu, E. Wang, Potential landscape and flux framework of nonequilibrium networks: robustness, dissipation, and coherence of biochemical oscillations. Proceedings of the National Academy of Sciences 105, 12271–12276 (2008).

82. E. H. Davidson, D. H. Erwin, Gene regulatory networks and the evolution of animal body plans. Science 311, 796–800 (2006).

83. N. G. Van Kampen, Stochastic processes in physics and chemistry. (Elsevier, 1992), vol. 1.

84. W. Zheng, H. Zhao, E. Mancera, L. M. Steinmetz, M. Snyder, Genetic analysis of variation in transcription factor binding in yeast. Nature 464, 1187–1191 (2010).

85. A. Anders, B. Ghosh, T. Glatter, V. Sourjik, Design of a MAPK signalling cascade balances energetic cost versus accuracy of information transmission. Nature communications 11, 1–10 (2020).

86. H. G. Dohlman, J. Song, D. Ma, W. E. Courchesne, J. Thorner, Sst2, a negative regulator of pheromone signaling in the yeast Saccharomyces cerevisiae: expression, localization, and genetic interaction and physical association with Gpa1 (the G-protein alpha subunit). Molecular and cellular biology 16, 5194–5209 (1996).

87. M. Sugiyama, Y. Kaneko, S. Harashima, Yeast protein phosphatases Ptp2p and Msg5p are involved in G1–S transition, CLN2 transcription, and vacuole morphogenesis. Archives of microbiology 191, 721-733 (2009).

88. W. A. Laviña, M. Sugiyama, Y. Kaneko, S. Harashima, Identification of protein kinase disruptions as suppressors of the calcium sensitivity of S. cerevisiae Δ ptp2 Δ msg5 protein phosphatase double disruptant. Archives of microbiology 192, 157–165 (2010).

89. W. A. Laviña, M. Sugiyama, Y. Kaneko, S. Harashima, Functionally redundant protein phosphatase genes PTP2 and MSG5 co-regulate the calcium signaling pathway in Saccharomyces cerevisiae upon exposure to high extracellular calcium concentration. Journal of bioscience and bioengineering 115, 138–146 (2013).

90. A.-C. Butty, P. M. Pryciak, L. S. Huang, I. Herskowitz, M. Peter, The role of Far1p in linking the heterotrimeric G protein to polarity establishment proteins during yeast mating. Science 282, 1511–1516 (1998).

91. A. Nern, R. A. Arkowitz, A Cdc24p-Far1p-Gβγ protein complex required for yeast orientation during mating. The Journal of cell biology 144, 1187–1202 (1999).

92. A. Nern, R. A. Arkowitz, Nucleocytoplasmic shuttling of the Cdc42p exchange factor Cdc24p. The Journal of cell biology 148, 1115–1122 (2000).

93. A. Nern, R. A. Arkowitz, A Cdc24p-Far1p-Gbetagamma protein complex required for yeast orientation during mating. J Cell Biol 144, 1187–1202 (1999).

94. E. H. Chen, Cell fusion: overviews and methods. (Springer, 2008).

95. E. Szczurek, I. GatLJViks, J. Tiuryn, M. Vingron, Elucidating regulatory mechanisms downstream of a signaling pathway using informative experiments. Molecular Systems Biology 5, 287 (2009).

96. C. I. Maeder et al., Spatial regulation of Fus3 MAP kinase activity through a reaction-diffusion mechanism in yeast pheromone signalling. Nature cell biology 9, 1319–1326 (2007).

97. A. Serrano et al., Spatio-temporal MAPK dynamics mediate cell behavior coordination during fungal somatic cell fusion. Journal of cell science 131, (2018).

98. J. E. Kranz, B. Satterberg, E. A. Elion, The MAP kinase Fus3 associates with and phosphorylates the upstream signaling component Ste5. Genes & Development 8, 313–327 (1994).

99. M. J. Winters, R. E. Lamson, H. Nakanishi, A. M. Neiman, P. M. Pryciak, A membrane binding domain in the Ste5 scaffold synergizes with Gβγ binding to control localization and signaling in pheromone response. Molecular cell 20, 21–32 (2005).

100. R. P. Bhattacharyya et al., The Ste5 scaffold allosterically modulates signaling output of the yeast mating pathway. Science 311, 822–826 (2006).

101. M. K. Malleshaiah, V. Shahrezaei, P. S. Swain, S. W. Michnick, The scaffold protein Ste5 directly controls a switch-like mating decision in yeast. Nature 465, 101–105 (2010).

102. T. Chen, J. Kurjan, Saccharomyces cerevisiae Mpt5p interacts with Sst2p and plays roles in pheromone sensitivity and recovery from pheromone arrest. Molecular and Cellular Biology 17, 3429–3439 (1997).

103. T. R. Garrison et al., Feedback phosphorylation of an RGS protein by MAP kinase in yeast. Journal of Biological Chemistry 274, 36387–36391 (1999).

104. S. C. Parnell et al., Phosphorylation of the RGS protein Sst2 by the MAP kinase Fus3 and use of Sst2 as a model to analyze determinants of substrate sequence specificity. Biochemistry 44, 8159–8166 (2005).

105. R. E. Chen, J. Thorner, Function and regulation in MAPK signaling pathways: lessons learned from the yeast Saccharomyces cerevisiae. Biochimica et Biophysica Acta (BBA)-Molecular Cell Research 1773, 1311–1340 (2007).

106. H.-O. Park, E. Bi, Central roles of small GTPases in the development of cell polarity in yeast and beyond. Microbiology and molecular biology reviews 71, 48–96 (2007).

107. J. B. Moseley, B. L. Goode, The yeast actin cytoskeleton: from cellular function to biochemical mechanism. Microbiology and Molecular Biology Reviews 70, 605–645 (2006).

108. A. Doncic, M. Falleur-Fettig, J. M. Skotheim, Distinct interactions select and maintain a specific cell fate. Molecular cell 43, 528–539 (2011).

109. N. Barkai, M. D. Rose, N. S. Wingreen, Protease helps yeast find mating partners. Nature 396, 422–423 (1998).

110. B. D. Slaughter, J. W. Schwartz, R. Li, Mapping dynamic protein interactions in MAP kinase signaling using live-cell fluorescence fluctuation spectroscopy and imaging. Proceedings of the National Academy of Sciences 104, 20320–20325 (2007).

111. E. A. Elion, M. Qi, W. Chen, Signaling specificity in yeast. Science 307, 687–688 (2005).

112. D. Fiedler et al., Functional organization of the S. cerevisiae phosphorylation network. Cell 136, 952–963 (2009).

113. J. Wang, Landscape and flux theory of non-equilibrium dynamical systems with application to biology. Advances in Physics 64, 1–137 (2015).

114. X. Fang, K. Kruse, T. Lu, J. Wang, Nonequilibrium physics in biology. Reviews of Modern Physics 91, 045004 (2019).

115. H. Feng, K. Zhang, J. Wang, Non-equilibrium transition state rate theory. Chemical Science 5, 3761–3769 (2014).

116. X. N. Fang, K. Kruse, T. Lu, J. Wang, Nonequilibrium physics in biology. Reviews of Modern Physics 91, (2019).

117. A. Nern, R. A. Arkowitz, G proteins mediate changes in cell shape by stabilizing the axis of polarity. Molecular cell 5, 853–864 (2000).

118. D. T. Gillespie, The chemical Langevin equation. The Journal of Chemical Physics 113, 297–306 (2000).

119. D. T. Gillespie, Approximate accelerated stochastic simulation of chemically reacting systems. The Journal of chemical physics 115, 1716–1733 (2001).

120. H. De Jong, Modeling and simulation of genetic regulatory systems: a literature review. Journal of computational biology 9, 67–103 (2002).

121. J. Lee Rodgers, W. A. Nicewander, Thirteen ways to look at the correlation coefficient. The American Statistician 42, 59–66 (1988).

122. A. Treuner-Lange, L. Sogaard-Andersen, Regulation of cell polarity in bacteria. J Cell Biol 206, 7–17 (2014).

123. J. Dworkin, Cellular Polarity in Prokaryotic Organisms. Csh Perspect Biol 1, (2009).

124. R. M. Macnab, D. E. Koshland, The gradient-sensing mechanism in bacterial chemotaxis. Proceedings of the National Academy of Sciences 69, 2509–2512 (1972).

125. L. Bardwell, A walk-through of the yeast mating pheromone response pathway. Peptides 25, 1465–1476 (2004).

126. S. Ghaemmaghami et al., Global analysis of protein expression in yeast. Nature 425, 737–741 (2003).

