## Supplementary Materials for "Global quantitative understanding of nonequilibrium cell fate decision making in response to pheromone"

**This PDF file includes:**

Materials and Methods

Figs. S1 to S14

Tables. S1 to S9

References

**Other Supplementary Materials for this manuscript include the following:**

Movies S1 to S9

**Materials and Methods**

**Pheromone environment**

The stimulus source we chose is pheromone, an alpha factor peptide hormone, whose molecular weight is 1683.98 (GenScript). We weigh a certain amount of powdered pheromone and dissolve them in YPD and YNB liquid medium respectively to prepare 1000μM pheromone medium, and then gradually dilute it to the ideal concentration medium for microscopy experiments and other experiments.

At the beginning, in order to adapt to the microfluidic environment, the yeast cells are cultured in the YNB conventional medium for 60 minutes, and then are switched to the medium supplemented with pheromone for culture.

**Dual fluorescence system construction**

When designing the vector for the gene editing of *FUS3*, we consulted the sequence of the plasmid “pFA6a-link-yomCherry-CaURA3”. This plasmid’s core sequence (“yomCherry-CaURA3”) was incorporated into the homologous recombination target fragment that we designed **(Fig. S10)**. The recombinant vector that we designed consists primarily of the left and right homology arms (F/R) of the gene *FUS3*, the flexible chain connecting *FUS3* and the fluorescent group (linker), the red fluorescent gene (yomCherry), and the screening gene (CaURA3). Among them, the “yomCherry” is a red fluorescent protein; the “CaURA3” is a protein that can compensate for the uracil deficiency-induced lethality of yeast.

First, 300bp were chosen as the left and right homologous long arms (F'/R') before and after the stop codon (TAG) of the *FUS3* gene. The homology arms (F/R) within the sequence interval of F'/R' can be obtained by performing PCR with these homologous long arm primers. Second, we selected a flexible chain (6aa [GS]x linker) as a structural buffer in order to fuse the red fluorescent reporter gene (“yomCherry”) to the 3' end of gene *FUS3* without destroying the protein structure between the two groups. Finally, the vector containing “yomCherry-CaURA3” was introduced into yeast cells. Table S9 lists the gene sequences we used.

**Real time image analysis**

Bright-field images obtained by the total internal reflection microscope were segmented, aligned, and labeled using a custom Matlab routine. We segmented the cells according to bright field images to obtain the outlines of the individual cells, and assigned each cell accordingly. Then we can collect the trajectories of the generations by the assigned id when the cells grow and divide, and obtain cell lineages. All the cell boundaries of yeast were manually corrected. The cell nuclei were distinguished by contouring the fluorescence images. Fluorescence intensity is the average of all fluorescence intensity within the cell boundary.

The real-time trajectories were obtained by automatic tracking, based on the cell overlaps between the adjacent frames. All trajectories require manual correction.

**Steady-state image analysis**

In order to explore the underlying mechanism of the bimodality, we collected the real time fluorescence intensity trajectories **(****Fig. 3A and Fig. S1–S2)**, which show that the yeast response is in a steady state after about 600min. This state means that the fluorescence intensity of the yeast inside and outside the nucleus does not increase or decrease significantly. The histograms of inner and outer fluorescence intensity were obtained by the steady-state fluorescence trajectories **(Fig. 3B and Fig. S3–S5)**.

The shape characteristics of the yeast are described by the $H_{n}$ in the yeast. The growth rate of the yeast is the numerical differentiation of the $H_{n}$. All the trajectories can be used to provide the quantitative analysis through Hidden Markov Chain Model (HMM). During the HMM fitting, the parameters of the fluorescence state were fixed. The distribution of high fluorescence state and low fluorescence state in two growth rates can be obtained by counting the state points on the trajectories.

**Data Analysis of the Time-Lapse Experiments**

The cell state can be distinguished from the trajectories using hidden Markov model (HMM). A maximum likelihood estimate of HMM parameters was performed globally on all the time traces separately. Multiple random initial parameters were used to start the iterative HMM analysis and ensure convergence to the global minimum. The Baum-Welch algorithm was used to re-estimate the parameters at the end of each iteration. Steady-state condition was enforced on the re-estimated parameters in each iteration. HMM can be directly applied to two-dimensional and high-dimensional data. Therefore, the inner and outer fluorescence data can be trained by the HMM, the results showed that the yeast fluorescence has bimodal distribution. The detailed results are shown in **Fig. 2**. The overlap ratio between the two states is obtained by integrating the overlapping part of the peak of the two states, which needs the results fitted by HMM. For each trajectory, the state can be determined by the HMM, and the state sequence is shown in **Fig. 3E**. The total residence time and the number of state changes can be obtained by counting the state points on the trajectories, and the average residence time is the quotient of the total residence time and the number of state changes.

**Distinguishing the boundaries of the nucleus**

When yeast cells are stimulated by pheromones, two responses occur: first, the cell cycle is arrested in the G1 phase, and second, mating-related genes are activated. To ensure both responses, large amounts of Fus3 are delivered to the nucleus, causing Fus3 to form fluorescent clusters in the nucleus (*1-9*). Previous articles have employed the fluorescent cluster region as the nucleus (*10-12*). When we observed cell budding and division under the microscope, the fluorescence clusters split in half, which corresponds to the separation of the daughter cell’s nucleus from the mother cell’s nucleus, indicating that it is feasible to use the fluorescence cluster as the nucleus. We use an algorithm similar to contour lines to divide each pixel in a cell into four distinct calculation levels. Then, the fluorescent spots at the same contour level are joined into closed regions. By comparing each level individually, it was determined that the outer edge of the highest level, which serves as the boundary between the nucleus and the cytoplasm, is best able to encompass intracellular fluorescent clusters **(Fig. S11).** Therefore, we propose that fluorescence above this threshold is localized within the nucleus, whereas fluorescence below this threshold is localized outside of the nucleus.

**Filled circle model in cell shape**

After identifying the boundary of the cell shape with “Matlab”, we filled it in with circles along the cell’s long axis in turns, with the area of the circle filled in each time being guaranteed to be the largest area of the remaining unfilled part. To ensure the accuracy of the cell morphology (H_n_) at various stages, we set a minimum circle diameter (5 pixels) to avoid significantly increasing the shape’s internal gap.

**Characterizing the deformation behavior of yeast cells**

To quantify the behavior of yeast cells of which deformation rate did not vary uniformly over time, we characterized various cell components using a circular fill pattern **(Fig. 5A)**. The largest circle represents the initial main part of the cell, while the smallest circle represents the newly formed portion at the top of the cell. For example, there are two filled circles denoted by the letters $R_{0}$ and $R_{1}$ within the yeast cell. $R_{0}$ is the larger of the two circles, whereas $R_{1}$ is the smaller **(Fig. S12)**. If the filled circle $R_{1}$ of the cells grows uniformly in size from small to large (model-1: “$R_{1\_1}\to R_{1\_2}\to R_{1\_3}\to R_{1\_4}$”), then the ratios of the smallest circle to the largest circle in the cell will be uniformly distributed. If, on the other hand, the cell grows as in model-2 (“$R_{1\_1}\to R_{1\_2}\to R_{1\_3}\to R_{1\_3}\to R_{1\_4}$”) with the growth temporarily halted at $R_{1\_3}$, then $R_{1\_3}$ will be observed repeatedly, increasing the probability of the $R_{1\_3}/R_{0}$. When the size ratio of the smallest and largest circles was used as an observable value, we noticed that the statistical result was mostly around 0.27 at 0.7 μM **(Fig. S13)**. This clearly demonstrated that when the size ratio was 0.27, the cell was growing more slowly or was temporarily not growing, and thus there were more opportunities to observe this ratio distribution during cell polar growth. That is, the rate of deformation or the capacity for growth at various locations within a cell were not exactly identical.

To quantify the various cell deformations that occur during cell growth, we considered a value (H_n_) comparable to the harmonic mean to characterize the cell morphology, i.e., $H_{n}=n(\frac{1}{R_{1}}+\frac{1}{R_{2}}+...+\frac{1}{R_{n}})$. The significant advantage of H_n_ is that it is particularly sensitive to small morphological changes at various locations of the cell. For example, we set the radius of the filled circle in the cell as R_1_, R_2_, and R_3_ in advance. If the cell grew longitudinally (in length), a new filled circle (R_4_) was added to the cell; consequently, the value of H_n_ increased as the number of elements in parentheses increased from three to four. In contrast, as cells expanded laterally (in width), the values of H_n_ decreased regardless of whether the filled circle’s radius increased.

**Derivation of the transition rates**

The master equation can be written as

$\frac{d}{dt}\binom{P_{1}}{P_{2}}=\left( \begin{matrix} k_{11} & k_{12} \\ k_{21} & k_{22} \end{matrix} \right)\binom{P_{1}}{P_{2}}=\left( \begin{matrix} -a & b \\ a & -b \end{matrix} \right)\binom{P_{1}}{P_{2}}$.

Here P_1_ and P_2_ are the probabilities of the low expression state and high expression state, respectively, while $k_{ij}$ (i, j=1, 2) is the transition rate from $P_{i}$ to $P_{j}$. We can write down the solution as follows with the initial conditions $P_{1}\left( 0 \right)=1$, $P_{2}\left( 0 \right)=0$,

$P_{1}\left( t \right)=\frac{b+ae^{-(a+b)t}}{a+b}$; $P_{2}\left( t \right)=-\frac{a(-1+e^{-\left( a+b \right)t})}{a+b}$.

Then the transition probability between the low expression state and the low expression state is $P_{11}=P_{1}\left( \delta t \right)=\frac{b+ae^{-\left( a+b \right)\delta t}}{a+b}$, where$\delta t$ is the observational time window for each time, here $\delta t=10 min$. With the initial conditions $P_{1}\left( 0 \right)=0$, $P_{2}\left( 0 \right)=1$,

$P_{1}\left( t \right)=-\frac{b(-1+e^{-\left( a+b \right)t})}{a+b}$; $P_{2}\left( t \right)=\frac{a+be^{-(a+b)t}}{a+b}$.

Then the transition probability between the high expression state and the high expression state is $P_{22}=P_{2}\left( \delta t \right)=\frac{a+be^{-\left( a+b \right)\delta t}}{a+b}$, where$\delta t$ is the observation time window for each time, here $\delta t=10 min$.

According to the HMM analysis, when the pheromone dose is 0.7 μM, the transition matrix is $P=(\begin{matrix} P_{11} & P_{12} \\ P_{21} & P_{22} \end{matrix}$)=($\begin{matrix} 0.6900 & 0.3100 \\ 0.3711 & 0.6289 \end{matrix}$). So we can get the transition rate as: a= 0.052017 (1/min), b= 0.062270 (1/min).

**Decomposing of the flux in the cell morphology**

The kinetics of the cell morphological can be modeled by a four-state Markov process. The transition probability ($M_{ij}$) can be calculated by counting the number of transitions in the state trajectories. Therefore, the master equation can be shown as follows:

$\frac{{dP}_{i}}{dt}=\sum_{j} M_{ij}P_{i}$	**(Equation 1)**

Where $P_{i}$ represents the probability of state $i$, and the transition probability $M_{ij}$ represents the transition probability from state $i$ to state $j$. For steady state, we set the left term of the master equation (1) to zero, then we obtain the steady state solution $P_{i}^{S}$, which is the long time limit. The steady state flux between state $i$ and $j$ can be defined as: $F_{ij}=-M_{ij}P_{i}^{S}+M_{ji}P_{j}^{S}$. If the steady state of the system is in equilibrium state, the flux between any two nodes in the system is zero, that is the detailed balance condition. For the general biological system, it does not necessarily satisfy the detailed balance condition ($F_{ij}\neq0$), the system is in non-equilibrium steady state, there will be at least one net flux among states.

In order to study the non-equilibrium steady states, we can separate the dynamical process into two parts, one is the detailed balance part and the other is the detailed balance-breaking kinetic process. To describe the detailed balance breaking, we decompose the probability matrix. The component of the rate matrix $MP$ can be decomposed into two parts, one being the symmetric matrix and the other cycle matrix, that is

$MP=C+D$	**(Equation 2)**

Where $D$ is the symmetric matrix $D_{ij}=\left( M_{ij}P_{i}^{S}+M_{ji}P_{j}^{S} \right)/2$, $C$ is the asymmetric matrix $C_{ij}=\left( M_{ij}P_{i}^{S}-M_{ji}P_{j}^{S} \right)/2$. Since the system contains four states, the asymmetric flux is not unique and contains three net cycle fluxes. Meanwhile, in order to ensure the consistency of decomposition for various pheromone concentration systems, the given base sets of flux decomposition were selected. Here, the base sets we selected are “State_1_ –State_2_ –State_3_ –State_1_”, “State_1_ –State_4_ –State_3_ –State_1_”, “State_2_ –State_4_ –State_3_ –State_2_”, the corresponding net fluxes are $J_{1}$, $J_{2}$ and $J_{3}$. The asymmetric matrix $C$ can be rewritten as follows:

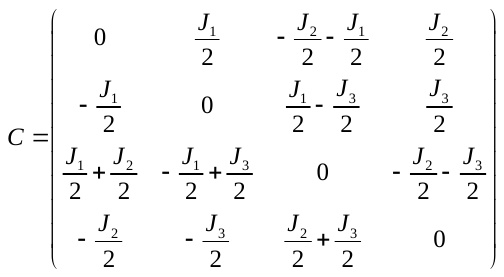

The linear equations can be obtained by corresponding the numerical results of the above matrix $C$ and the experimental statistics **(Table S1–S5)**. For example, three linearly independent parameters of $C_{1,4}, C_{1,3}, C_{2,3}$ are selected for the linear equations originated from the underlying master equations.

The linear equations (0.7 μM) are
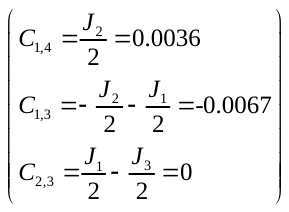
. Then, solving these linear equations one can get the net flux values of the non-equilibrium system.

**The comprehensive description of the biochemical reactions in stochastic simulations.**

The biochemical reactions in our signal transduction model were derived from the preexisting gene regulatory network. These biochemical reactions include the processes of gene translation into the proteins, the phosphorylation of the proteins, the degradation of the proteins, the interaction between the proteins, and the transfer of the proteins within the nucleus and the cytoplasm of the cells. The following is an understanding of each chemical reaction **(Table S8)**:

Reaction 1 depicts a portion of the gene *FUS3* being translated into a cytoplasmic protein at a certain rate. Reaction 2 depicts the gene *FUS3* generated by the indirect pheromone induction in addition to the known gene regulation. Reaction 3 is the reverse reaction of the reaction 2. Reaction 4 depicts the process of the phosphorylation of the protein produced by the gene *FUS3* induced by the pheromones outside the cell nucleus. Reaction 5 represents the gene *FUS3* produced by the self-activation of the phosphorylated protein Fus3 outside the cell nucleus. Reaction 6 is the reverse reaction of the reaction 5. Reaction 7 represents the process of the phosphorylation of a protein generated outside of the nucleus by the self-activating gene *FUS3*. Reaction 8 represents the transport of the phosphorylated Fus3 from the outside of the nucleus to the inside of the nucleus. Reaction 9 is the reverse reaction of the reaction 8.

Reactions 10 and 11 represent the degradation of the phosphorylated Fus3 both outside and inside the nucleus. Reaction 12 represents the translation of the gene *STE12* into the phosphorylated protein Ste12. Reaction 13 represents the impact of the gene *STE12* on the nuclear protein Fus3. Reaction 14 is the reverse reaction of the reaction 13. Reaction 15 represents the activation of the gene *STE12* by the nuclear Fus3. Reaction 16 represents the process of the translation of the gene *MSG5* into the phosphorylated protein Msg5. Reaction 17 represents the interaction between the phosphorylated protein Ste12 and the gene *MSG5*. Reaction 18 is the reverse reaction of the reaction 17. Reaction 19 represents the process of the translation of the gene *MSG5*, which is activated by the phosphorylated protein Ste12.

Reaction 20 represents the interaction between the phosphorylated protein *MSG5* and the gene *FUS3*. Reaction 21 is the reverse reaction of the reaction 20. The reaction 22 represents the inhibition of the gene *FUS3* expression by the phosphorylated protein *MSG5*. Reactions 23 and 24 represent the degradation of the proteins Ste12 and Msg5, respectively. Reaction 25 represents the process of the phosphorylation of protein Far1 generated by the gene *FAR1*. Reaction 26 represents the interaction between the Fus3 in the nucleus and the gene *FAR1*. Reaction 27 is the reverse reaction of reaction the 26. Reaction 28 represents the process of the transcription of the gene *FAR1*, regulated by the Fus3, into the protein. Reaction 29 represents the degradation of the protein Far1.

Reaction 30 represents the process of the gene *BNI1* generating the protein Bni1. Reaction 31 represents the interaction between the cytoplasmic Fus3 and the gene *BNI1*. Reaction 32 is the reverse reaction of the reaction 31. Reaction 33 represents the process of the indirect inhibition of the Fus3 expression outside of the nucleus by Bni1. Reaction 34 represents the interaction between the nuclear gene *BNI1* and the phosphorylated Far1. Reaction 35 is the reverse reaction of the reaction 34. Reaction 36 represents the process of the phosphorylated Far1-activated generation of the phosphorylated protein from the gene *BNI1*. Reaction 37 represents the interaction between the nuclear Fus3 and the gene *BNI1*. Reaction 38 is the reverse reaction of the reaction 37. Reaction 39 represents the process of the generation of phosphorylated protein from the gene *BNI1* activated by the phosphorylated Fus3 outside of the nucleus.

Reactions 40 and 41 denote the degradation of the phosphoproteins Bni1 both in the cytoplasmic and nuclear compartments.

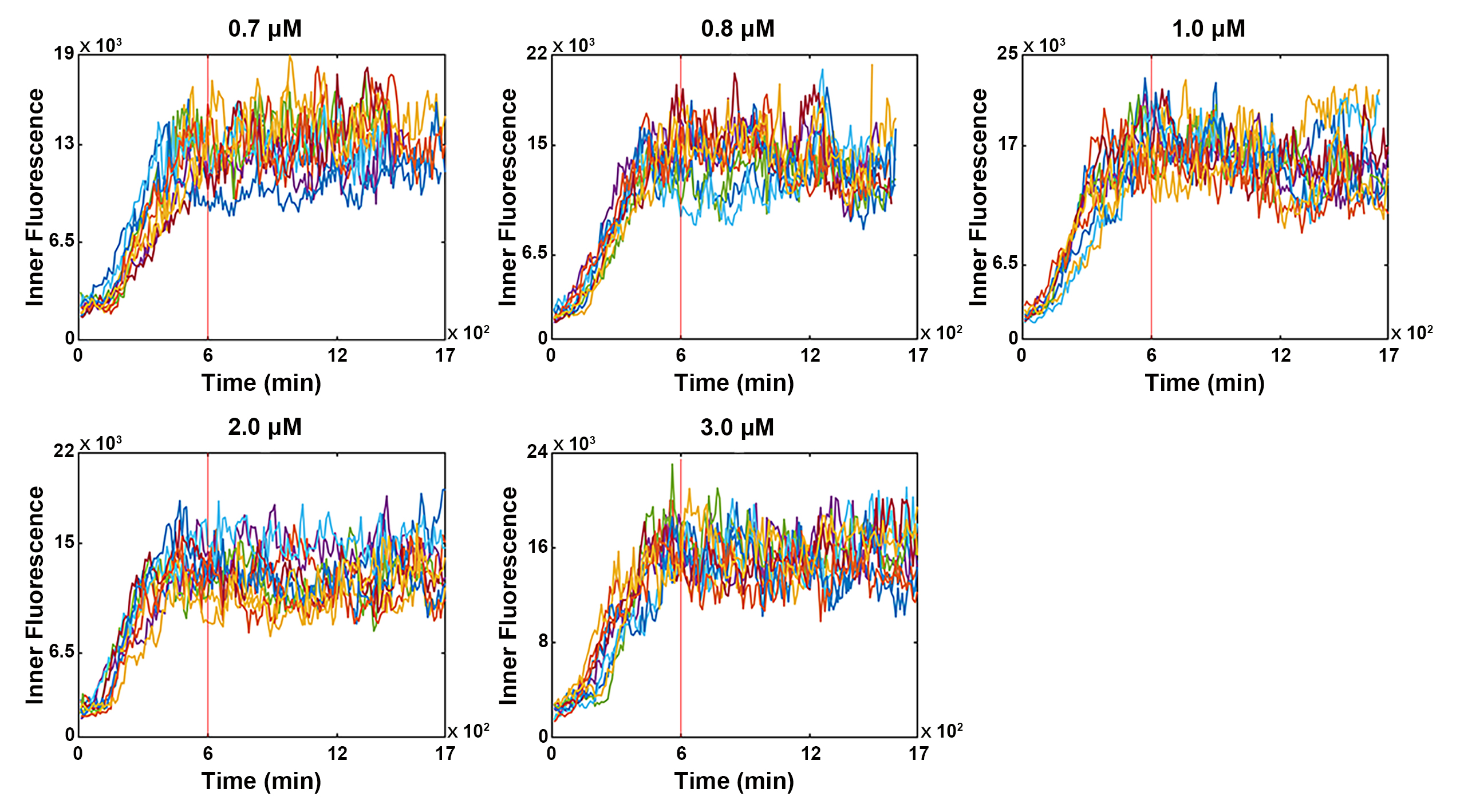

**Fig. S1. The Partial fluorescence intensity trajectories of Fus3 inside the nucleus under the different pheromone doses.** The red vertical line at 600 minutes is used to approximate the time node at which all cell fluorescence trajectories have entered a non-equilibrium steady state.

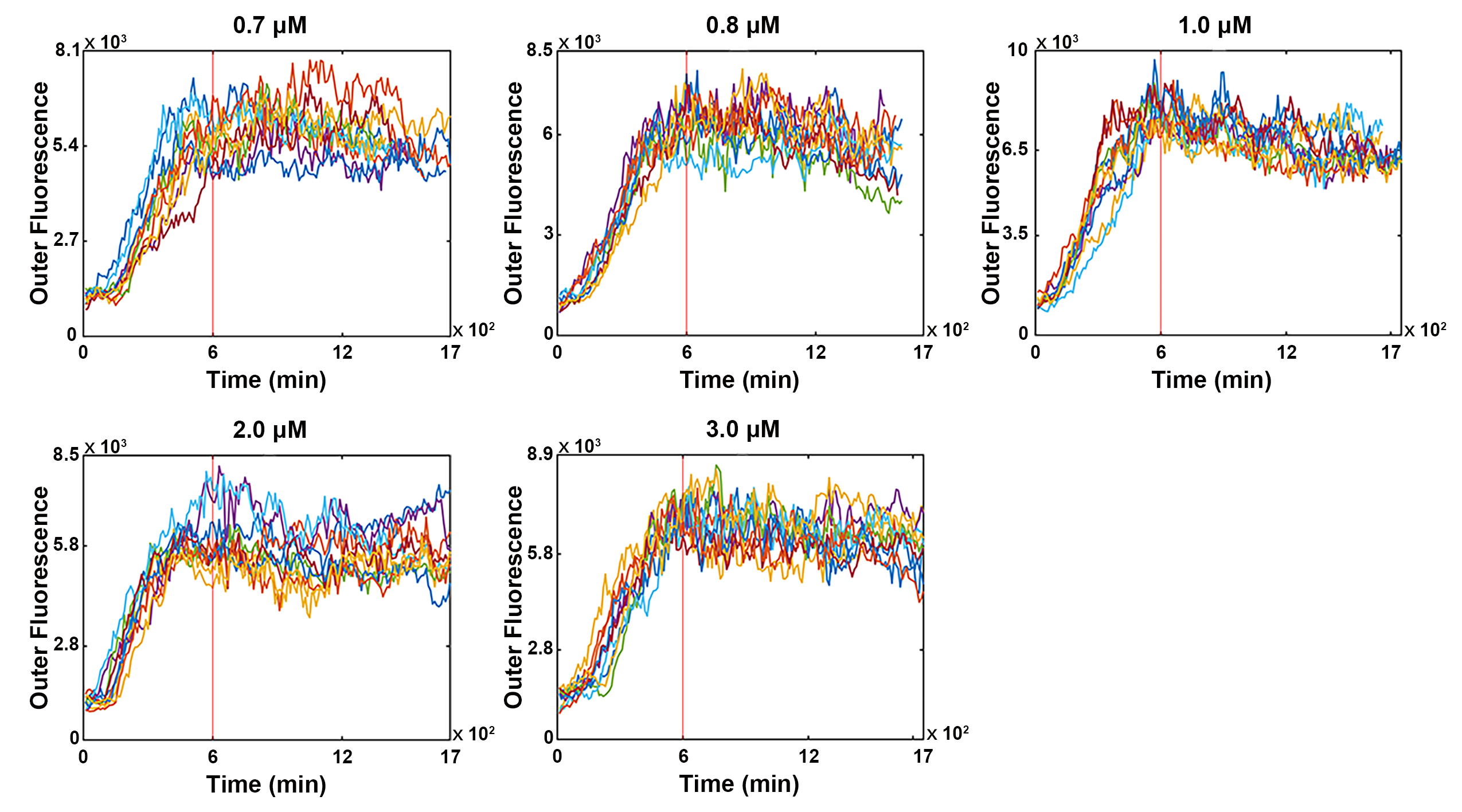

**Fig. S2. The Partial fluorescence intensity trajectories of Fus3 outside the nucleus under the different pheromone doses.** The red vertical line at 600 minutes is used to approximate the time node at which all cell fluorescence trajectories have entered a non-equilibrium steady state.

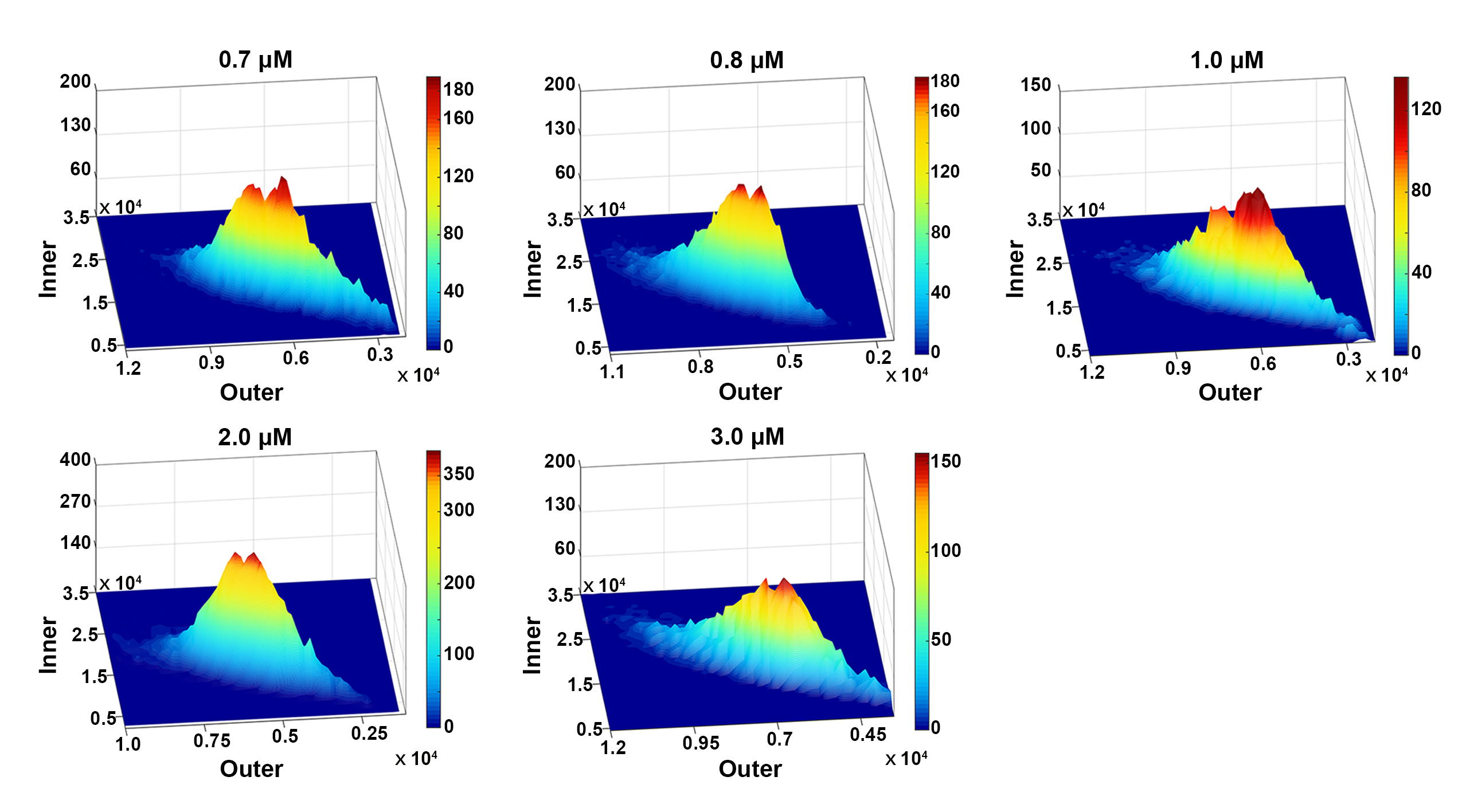
 **Fig. S3. The 3D distribution graph of Fus3 fluorescence intensity inside and outside the nucleus under the different pheromone doses.**

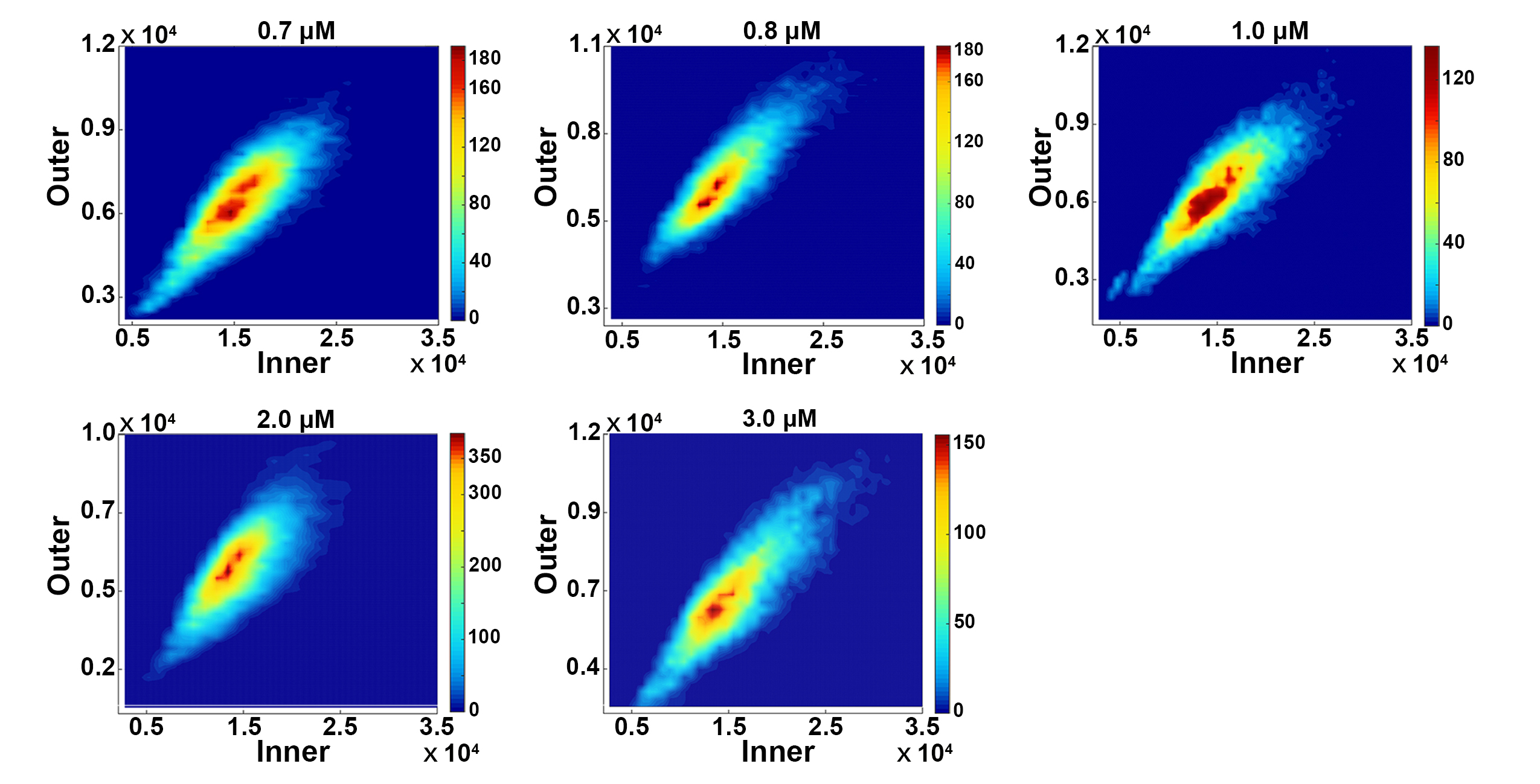
 **Fig. S4.** **The 2D distribution graph of Fus3 fluorescence intensity inside and outside the nucleus under the different pheromone doses.**

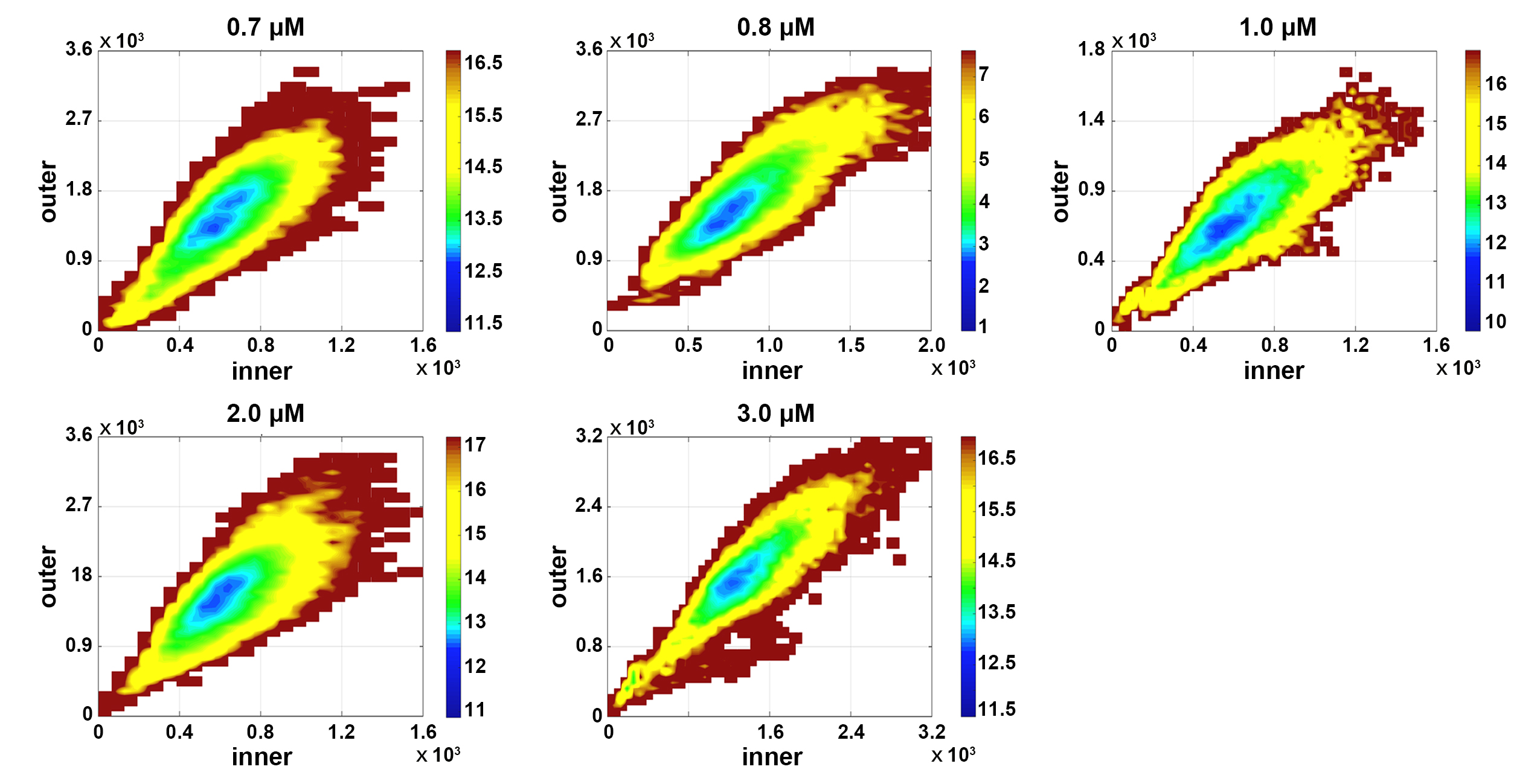
 **Fig. S5. The 2D underlying potential landscapes U under the different pheromone doses.**

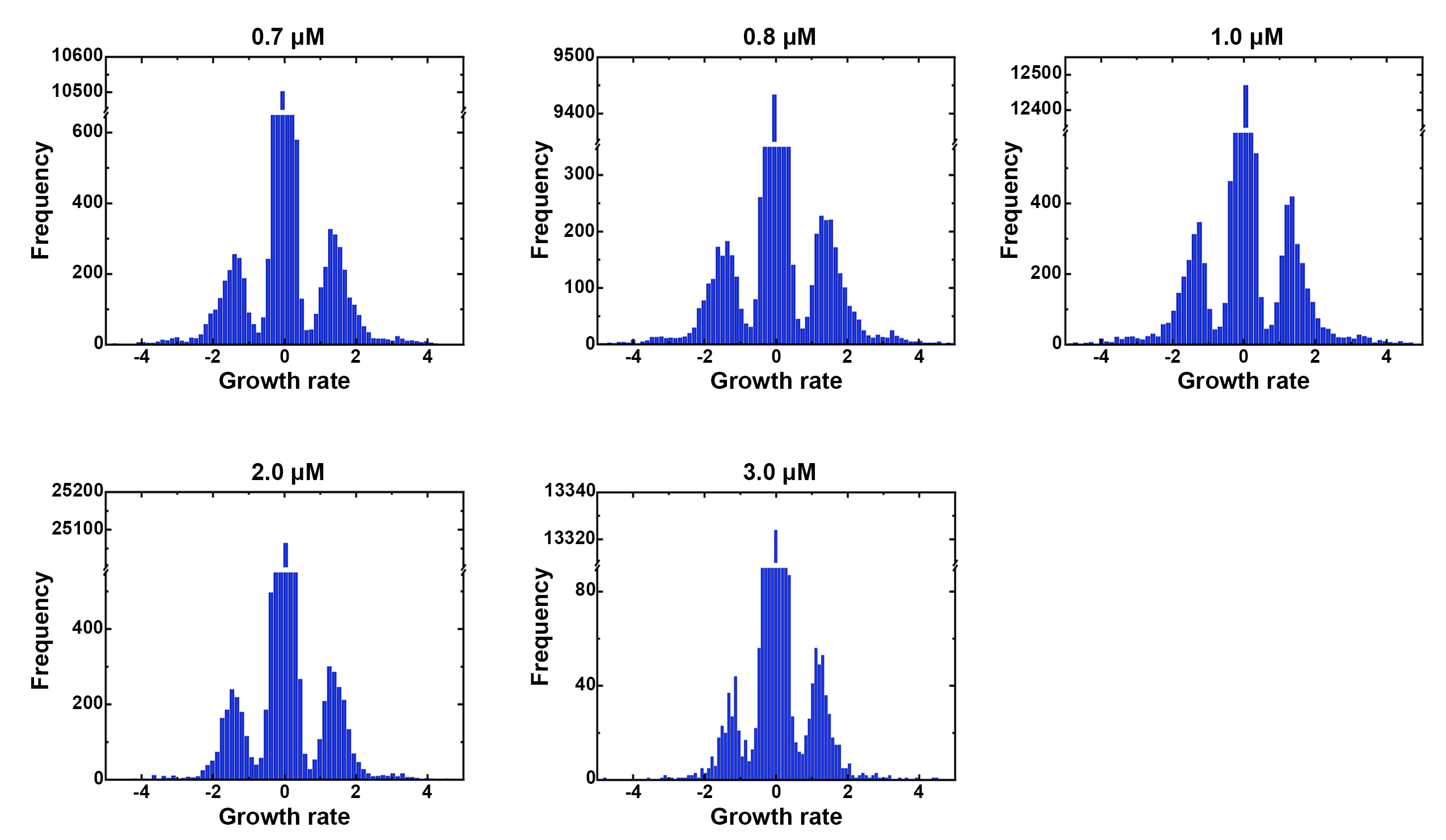
**Fig. S6. Statistical graph of the cell growth rate under the different pheromone doses.**

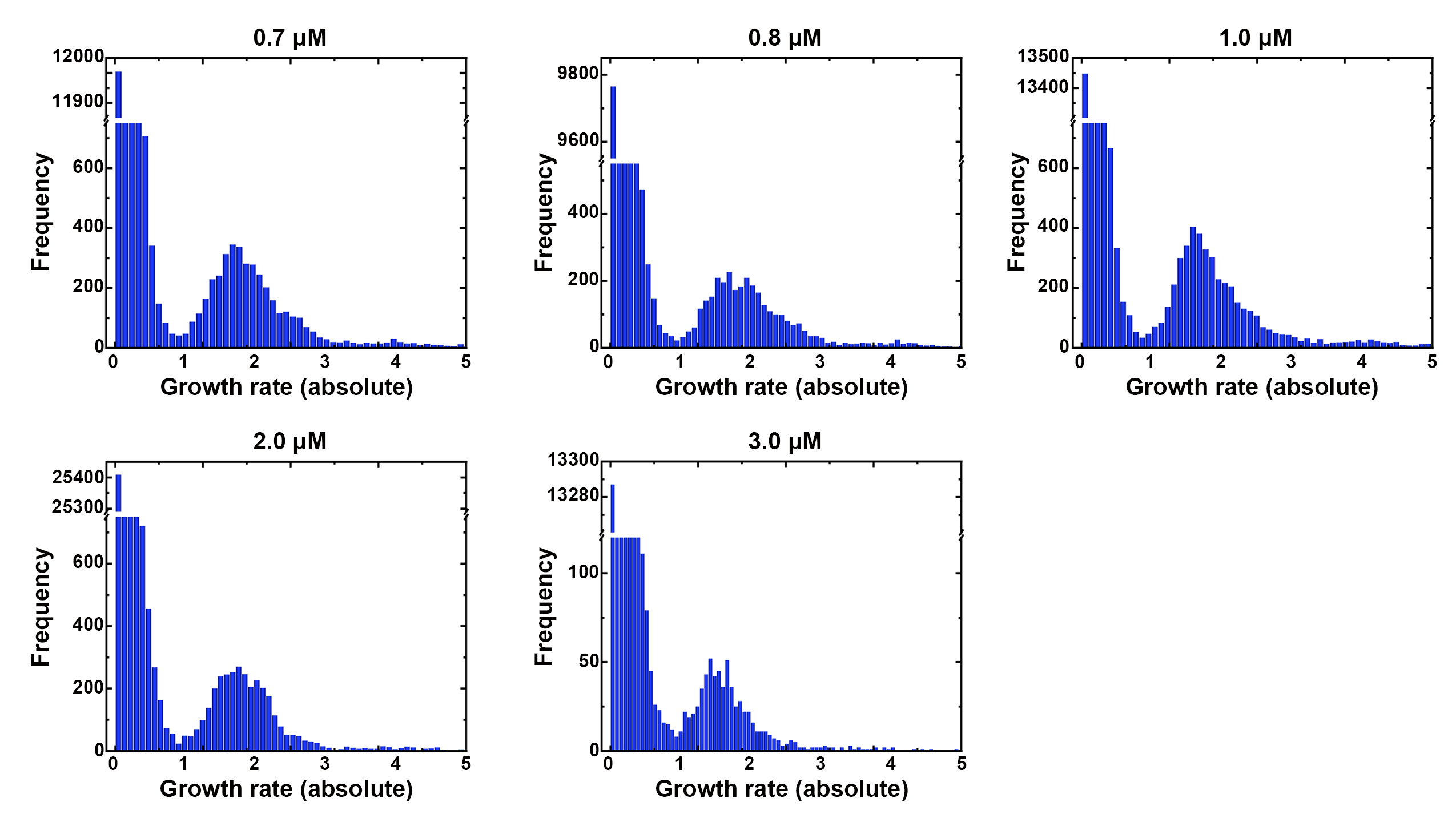
**Fig. S7. Statistical graph depicting the absolute value of the cell growth rate at various pheromone concentrations.**

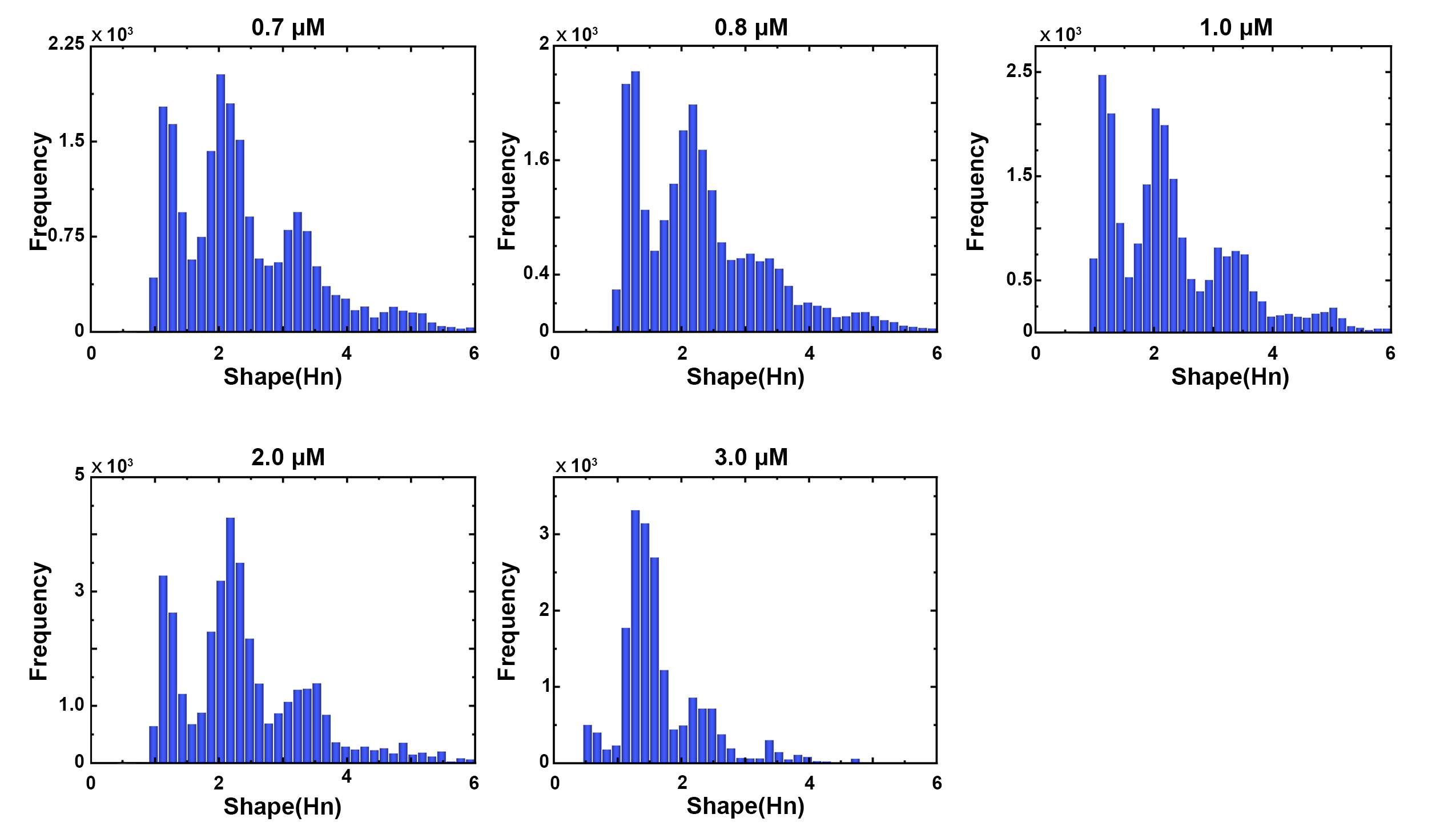
 **Fig. S8. The Statistical distribution of the cell morphological fates under the different pheromone doses.**

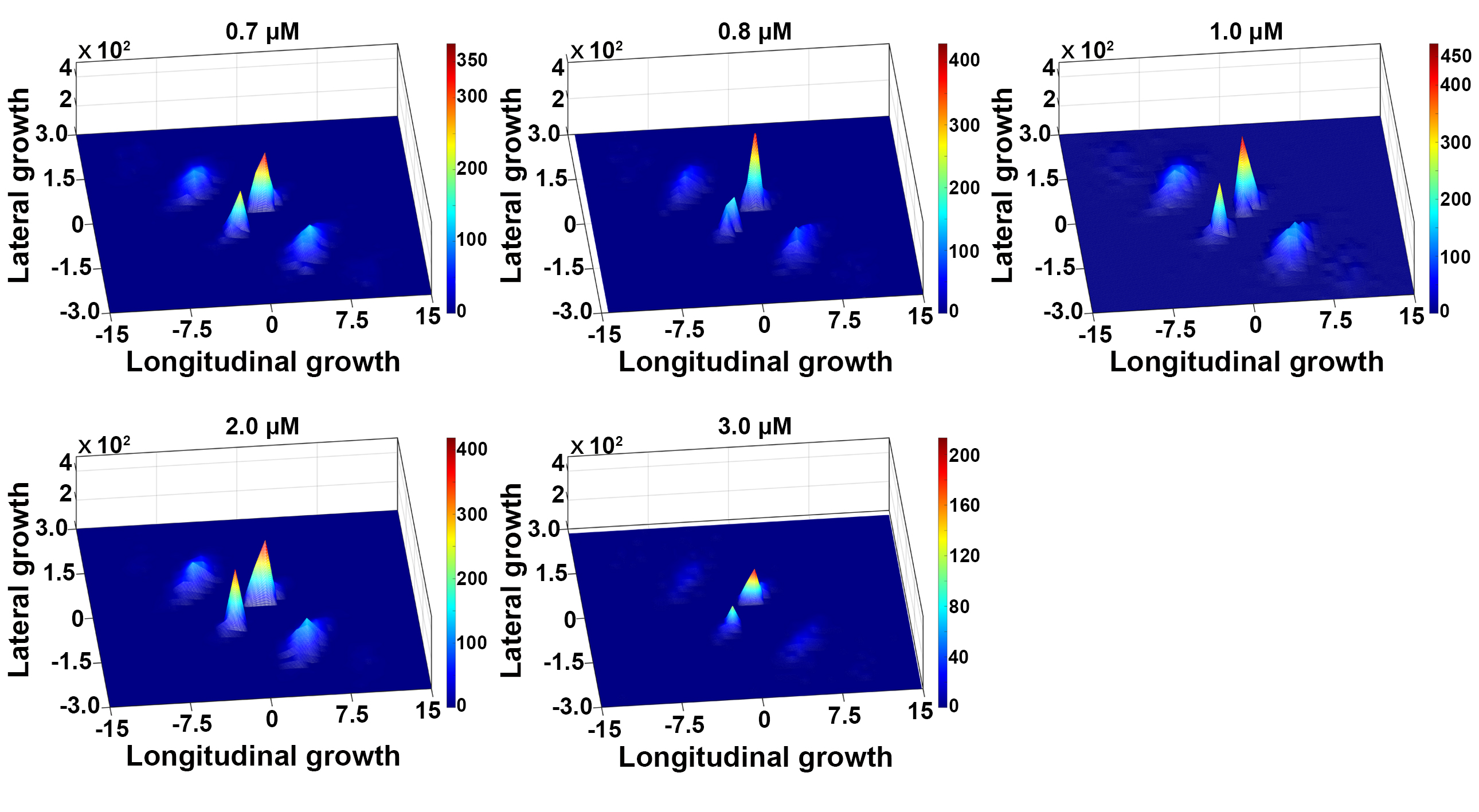

**Fig. S9. The synergistic effect of both lateral and longitudinal cell growth capabilities at various pheromone concentrations.** The x-axis represents the ability of cells to grow longitudinally, as indicated by the rate of change in the sum of the radii of the filled circles within the cells, i.e., ${{(R}_{1}+R_{2}+\ldots+R_{n})}^{'}$; the y-axis represents the ability of cells to grow laterally, as indicated by the rate of change in the average radius of the filled circles within the cells, i.e., ${(\frac{R_{1}+R_{2}+\ldots+R_{n}}{n})}^{'}$.

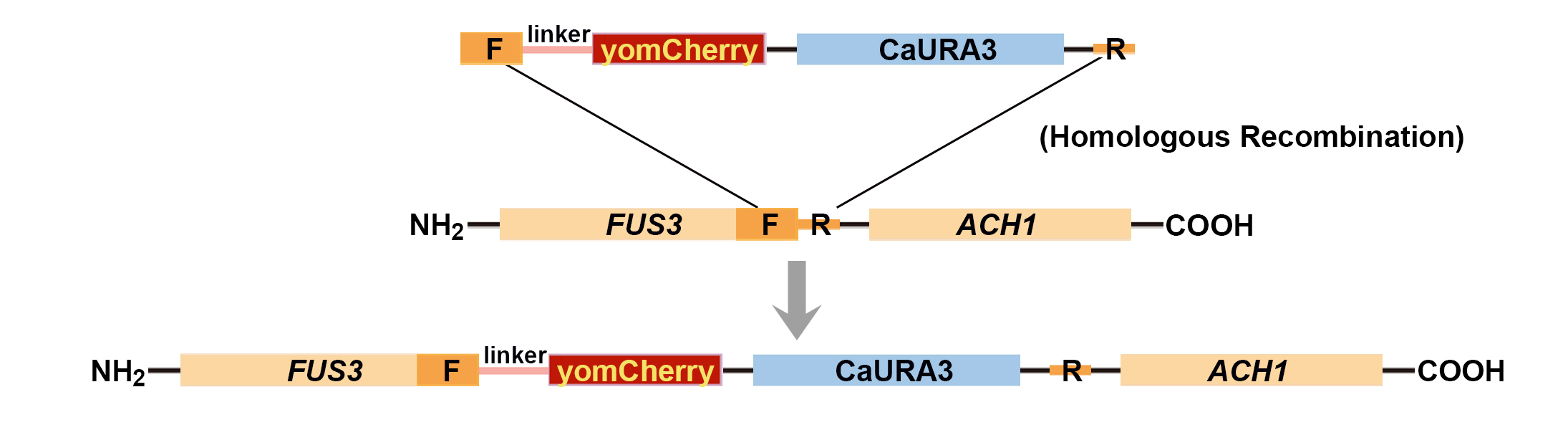

**Fig. S10. Diagram illustrating the homologous recombination principle of *FUS3*-yomCherry.** F/R is the left and right homology arms; the red fluorescent gene (yomCherry); the screening gene (CaURA3); the linker is a flexible chain (6aa [GS]x linker); Ach1 is the gene adjacent to the 3' end of *FUS3***.**

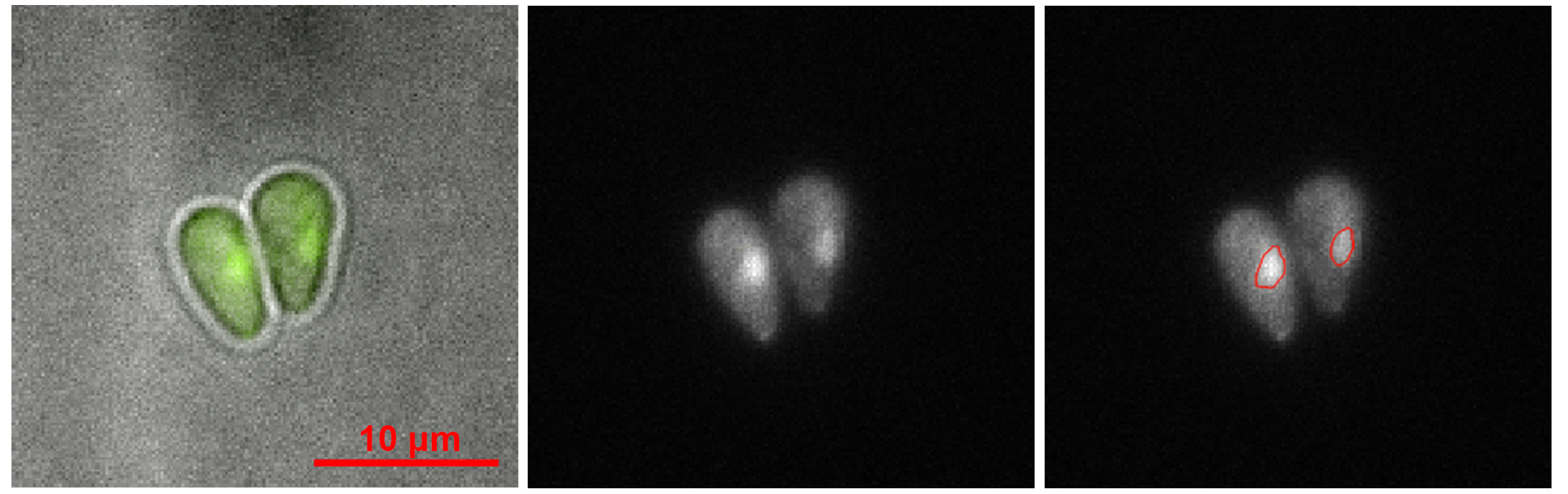

**Fig. S11. Delineation of the cell nucleus’s borders.** The cells depicted in the image are yeast cells incubated with 0.7 uM pheromone for 280min; the GFP (S65T) linked to *FUS3* was excited by a 488 nm laser; the image on the left is a superposition of the fluorescence field and bright field, allowing the outline of yeast cells to be seen; the brightness of the images in the middle and on the right represents the gene expression level of Fus3; the red circles in the right panel depict the calculated nucleus boundaries.

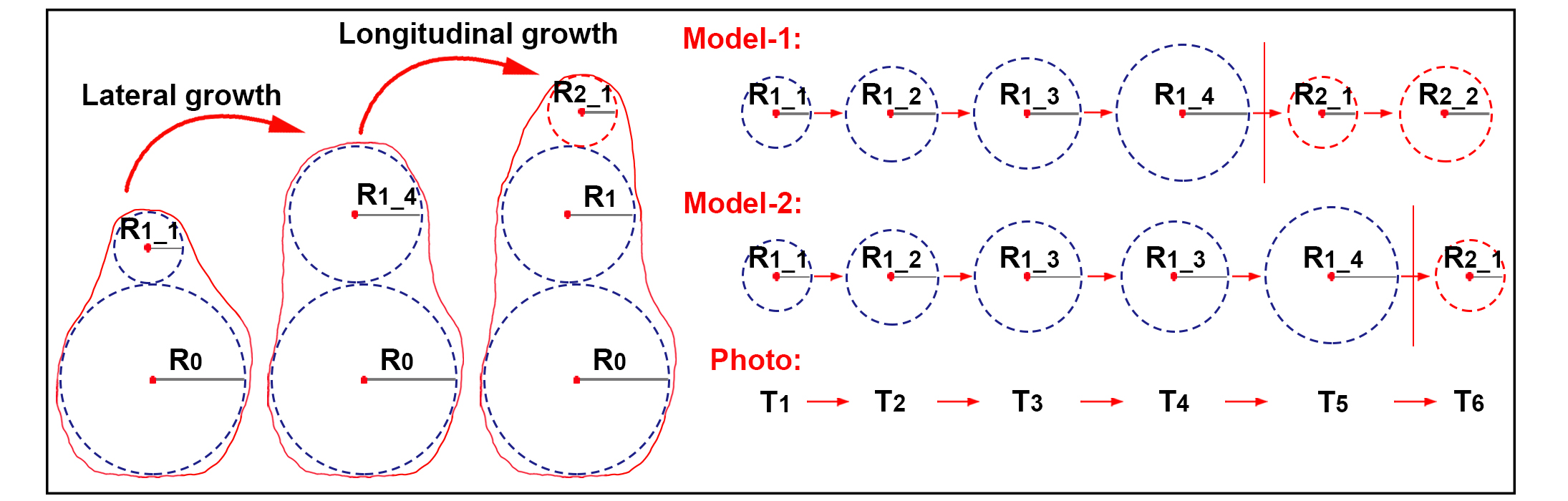

**Fig. S12. Schematic illustration of two models of cell growth.** $R_{0}$ is the maximum circle radius; $R_{1}$ is the minimum circle radius; the $(R_{1\_1}),...,(R_{1\_4})$ are the radii of $R_{1}$ in the different growth stages; $(T_{1},...,T_{6})$ are the times at which the microscope recorded the cell state.

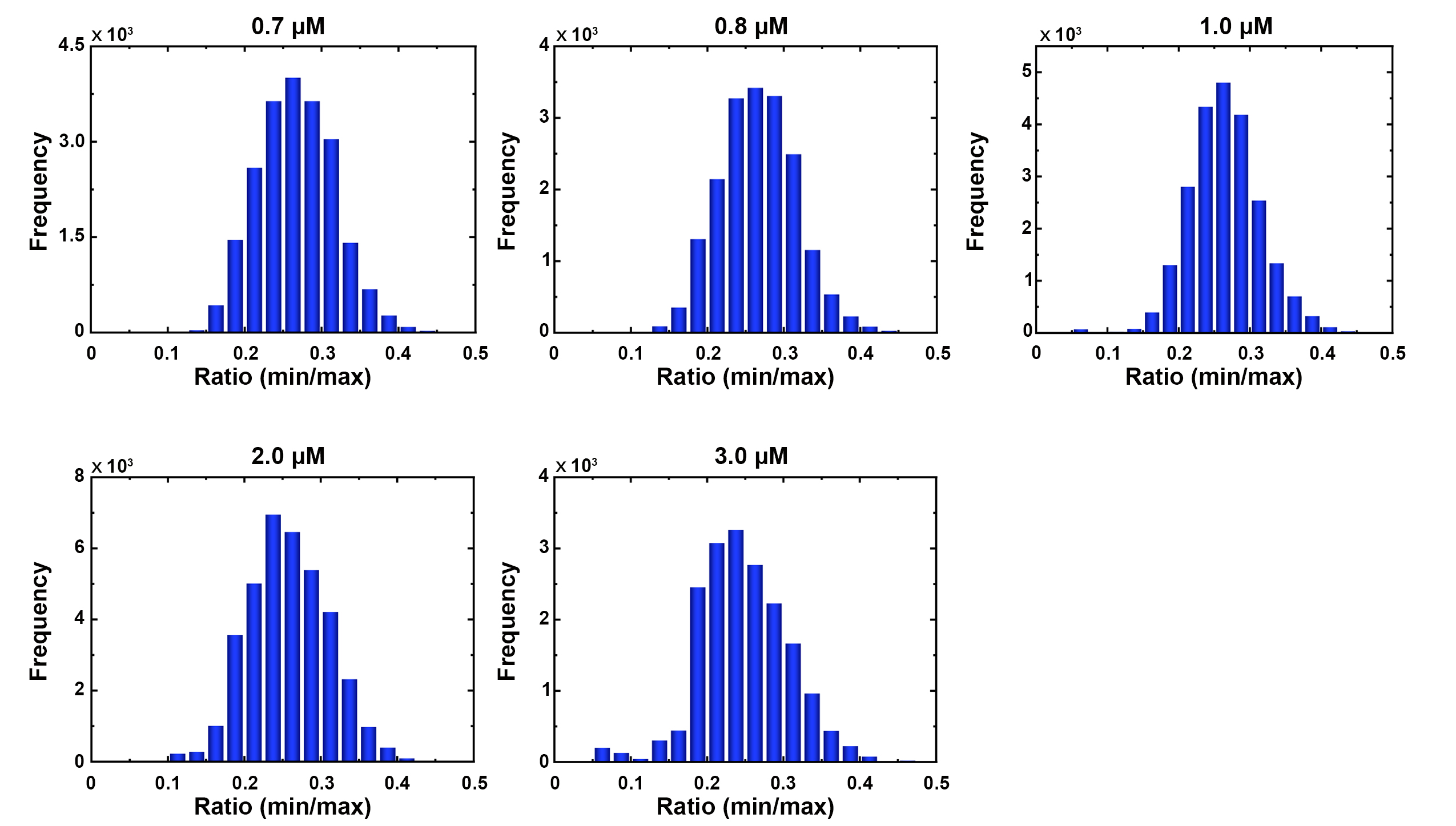
**Fig. S13. The distribution diagram of the radius ratio between the smallest circle and the largest circle of the yeast cell shape under the different pheromone doses.**

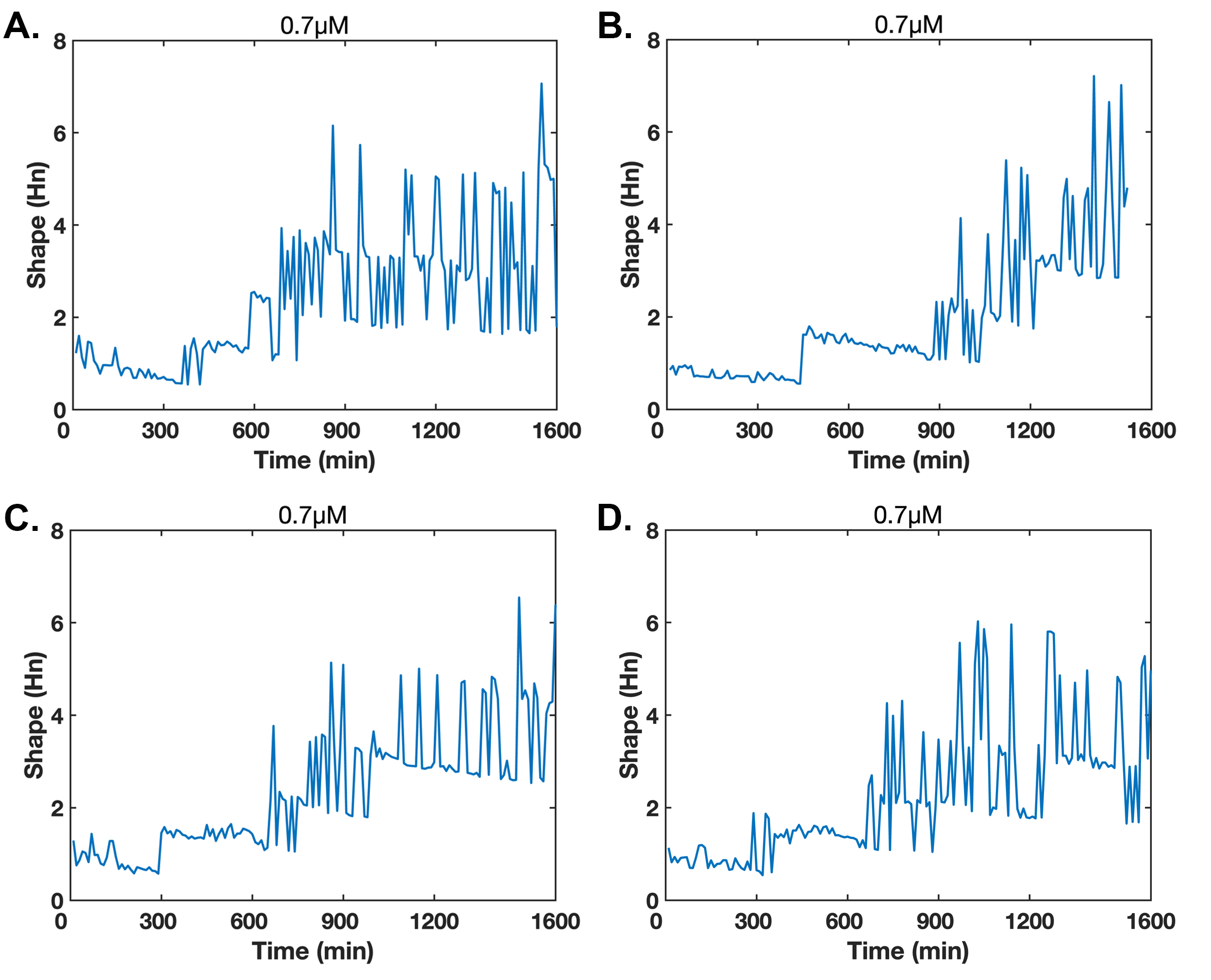

**Fig. S14. Real-time trajectory of the cell morphology (**$\mathbf{H}_{\mathbf{n}}$**) at 0.7 μM.**

**Table S1. The quantitative parameters in the net flux theory.** Following the master equation, parameters of net flux are determined by fitting the real-time trajectories of cell morphology at 0.7 μM to hidden Markov chain models.

| **Pheromone dose (μM)** | **0.7** |
| --- | --- |
| **Number of transitions** | 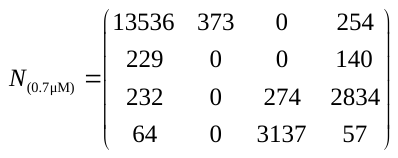 |
| **Transition probability** | 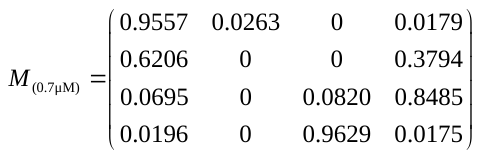 |
| **Probability of the four states** | 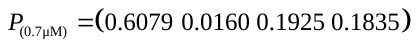 |
| **Asymmetric matrix** | 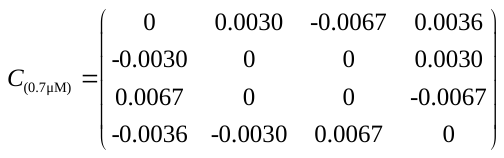 |
| **Symmetric matrix** | 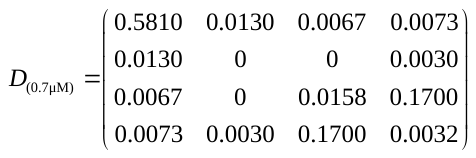 |
| **Net fluxes value** | 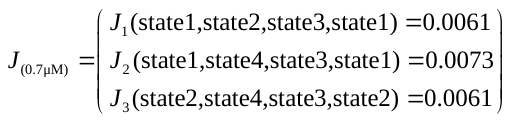 |

**Table S2. The quantitative parameters in the net flux theory.** Following the master equation, parameters of net flux are determined by fitting the real-time trajectories of cell morphology at 0.8 μM to hidden Markov chain models.

| **Pheromone dose (μM)** | **0.8** |
| --- | --- |
| **Number of transitions** | 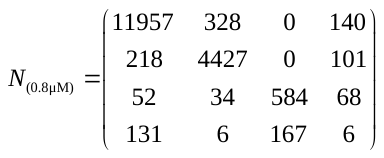 |
| **Transition probability** | 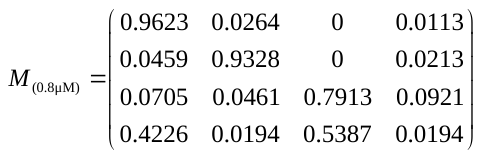 |
| **Probability of the four states** | 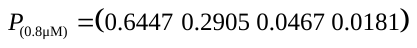 |
| **Asymmetric matrix** | 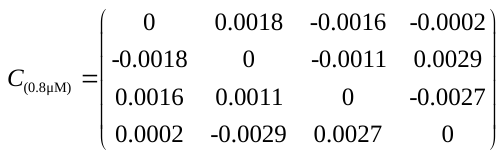 |
| **Symmetric matrix** | 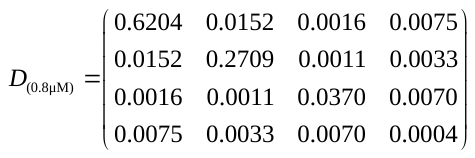 |
| **Net fluxes value** | 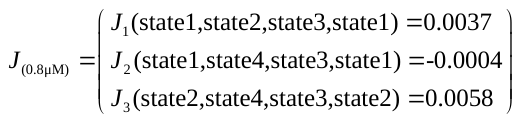 |

**Table S3. The quantitative parameters in the net flux theory.** Following the master equation, parameters of net flux are determined by fitting the real-time trajectories of cell morphology at 1.0 μM to hidden Markov chain models.

| **Pheromone dose (μM)** | **1.0** |
| --- | --- |
| **Number of transitions** | 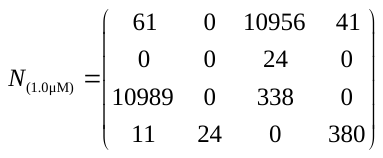 |
| **Transition probability** | 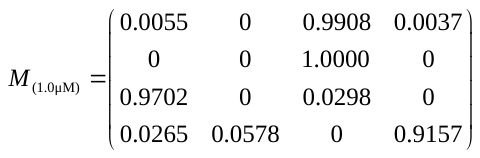 |
| **Probability of the four states** | 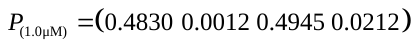 |
| **Asymmetric matrix** | 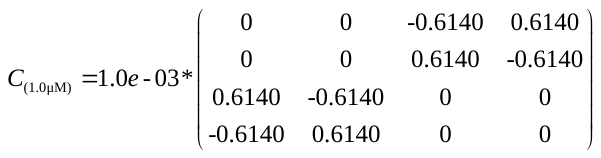 |
| **Symmetric matrix** | 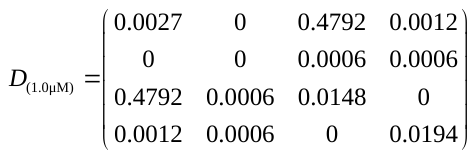 |
| **Net fluxes value** | 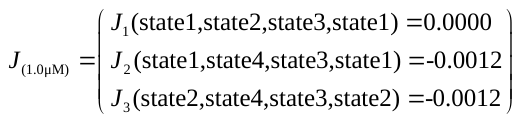 |

**Table S4. The quantitative parameters in the net flux theory.** Following the master equation, parameters of net flux are determined by fitting the real-time trajectories of cell morphology at 2.0 μM to hidden Markov chain models.

| **Pheromone dose (μM)** | **2.0** |
| --- | --- |
| **Number of transitions** | 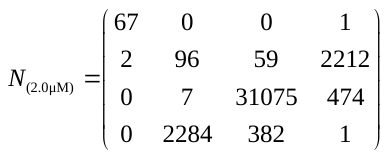 |
| **Transition probability** | 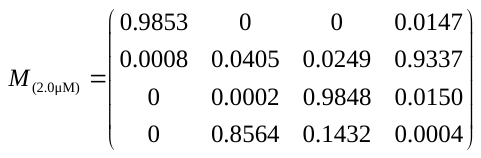 |
| **Probability of the four states** | 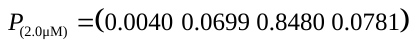 |
| **Asymmetric matrix** | 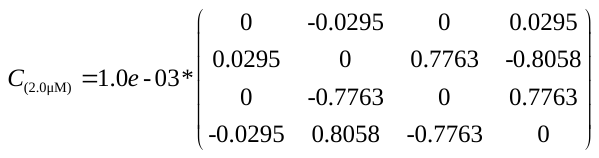 |
| **Symmetric matrix** | 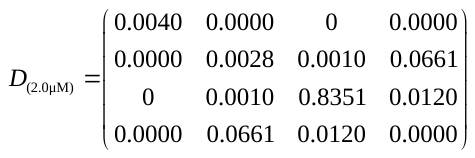 |
| **Net fluxes value** | 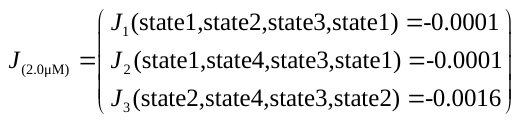 |

**Table S5. The quantitative parameters in the net flux theory.** Following the master equation, parameters of net flux are determined by fitting the real-time trajectories of cell morphology at 3.0 μM to hidden Markov chain models.

| **Pheromone dose (μM)** | **3.0** |
| --- | --- |
| **Number of transitions** | 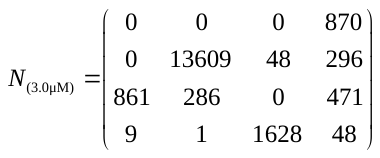 |
| **Transition probability** | 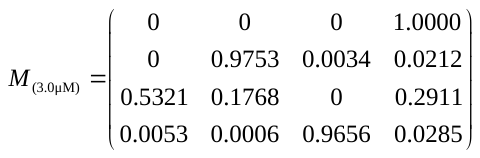 |
| **Probability of the four states** | 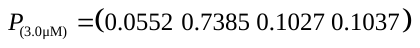 |
| **Asymmetric matrix** | 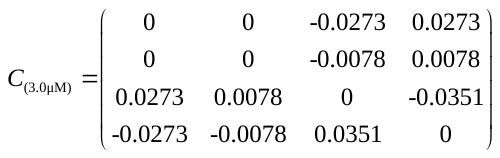 |
| **Symmetric matrix** | 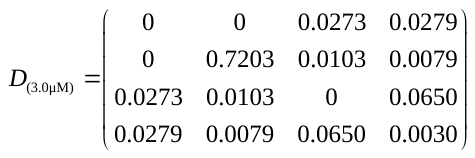 |
| **Net fluxes value** | 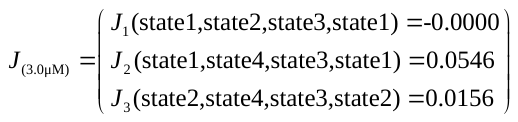 |

**Table S6. The description of the components in the biochemical reactions.**

| **Name** | **Description** |
| --- | --- |
| $GFus3$ or *FUS3* | The gene of the *FUS3* |
| $PFus3$ or Fus3 | The protein of the *FUS3* |
| $P{Fus3}_{p}$ or ${Fus3}_{p}$ | Phosphorylated protein of the *FUS3* |
| ${(P{Fus3}_{p})}^{'}$ or ${({Fus3}_{p})}^{'}$ | The $P{Fus3}_{p}$ remaining after the chemical reaction |
| $P{Fus3}_{p}\text{\_inner}$ | The $P{Fus3}_{p}$ in the nucleus |
| $P{Fus3}_{p}\text{\_outer}$ | The $P{Fus3}_{p}$ outside the nucleus |
| $P{Fus3}_{p}\text{\_in}$ | The $P{Fus3}_{p}$ produced by the $P_{1}$ pathway in the nucleus |
| $P{Fus3}_{p}\text{\_out}$ | The $P{Fus3}_{p}$ produced by the $P_{2}$ pathway outside the nucleus |
| $PFus3\text{-B}$ | This indicates the $P{Fus3}_{p}$ produced by the action of the B |
| $GFus3\text{-self}$ | The $GFus3$ produced by its self-activation |
| $\text{Φ}$ | Protein degradation |

**Table S7. The list of the components in the biochemical reactions.**

| **1** | 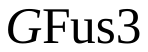 | **21** | 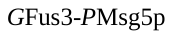 |
| --- | --- | --- | --- |
| **2** | 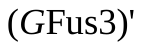 | **22** | 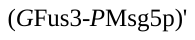 |
| **3** | 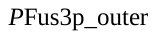 | **23** | 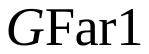 |
| **4** | 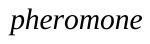 | **24** | 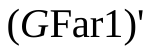 |
| **5** | 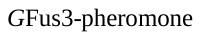 | **25** | 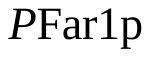 |
| **6** | 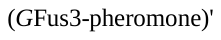 | **26** | 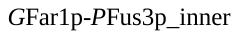 |
| **7** | 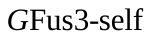 | **27** | 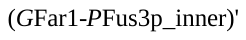 |
| **8** | 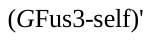 | **28** | 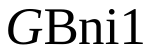 |
| **9** | 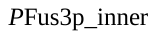 | **29** | 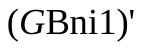 |
| **10** | 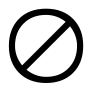 | **30** | 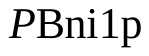 |
| **11** | 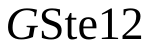 | **31** | 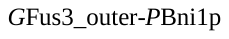 |
| **12** | 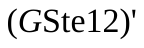 | **32** | 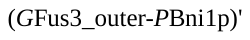 |
| **13** | 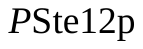 | **33** | 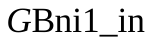 |
| **14** | 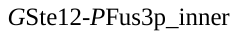 | **34** | 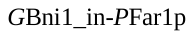 |
| **15** | 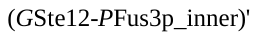 | **35** | 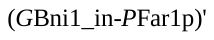 |
| **16** | 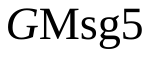 | **36** | 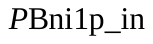 |
| **17** | 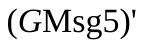 | **37** | 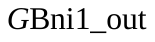 |
| **18** | 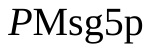 | **38** | 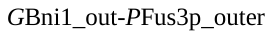 |
| **19** | 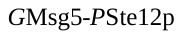 | **39** | 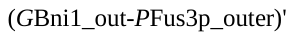 |
| **20** | 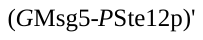 | **40** | 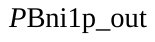 |

**Table S8. The list of the biochemical reactions.**

| **1** | 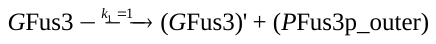 |
| --- | --- |
| **2** | 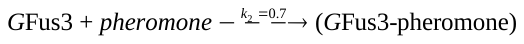 |
| **3** | 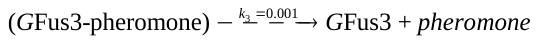 |
| **4** | 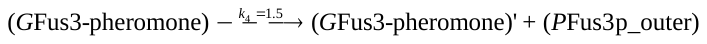 |
| **5** | 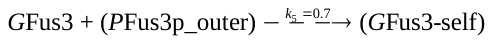 |
| **6** | 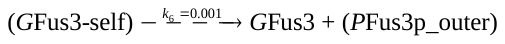 |
| **7** | 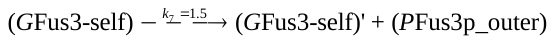 |
| **8** | 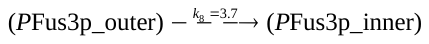 |
| **9** | 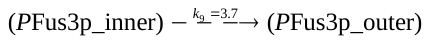 |
| **10** | 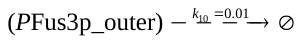 |
| **11** | 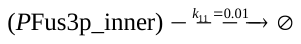 |
| **12** | 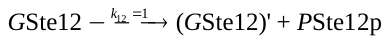 |
| **13** | 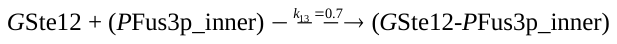 |
| **14** | 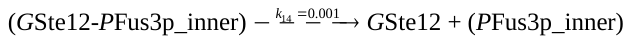 |
| **15** | 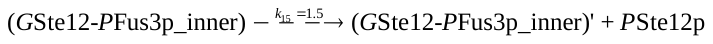 |
| **16** | 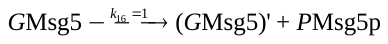 |
| **17** | 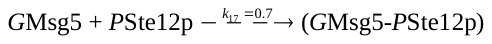 |
| **18** | 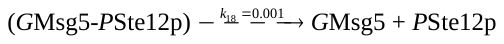 |
| **19** | 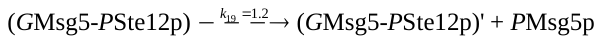 |
| **20** | 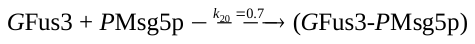 |
| **21** | 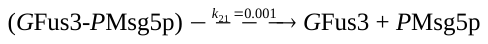 |
| **22** | 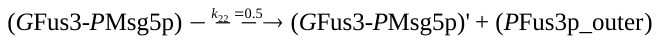 |
| **23** | 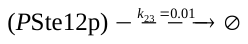 |
| **24** | 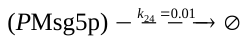 |
| **25** | 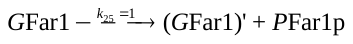 |
| **26** | 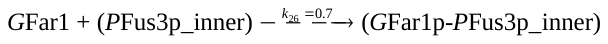 |
| **27** | 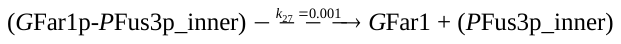 |
| **28** | 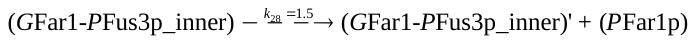 |
| **29** |  |
| **30** |  |
| **31** |  |
| **32** |  |
| **33** |  |
| **34** |  |
| **35** |  |
| **36** |  |
| **37** |  |
| **38** |  |
| **39** |  |
| **40** |  |
| **41** |  |

**Table S9. The components of the homologous recombination vector.**

| **Name** | **Sequence** |
| --- | --- |
| homologous long arms F' (300 bp**)** | CCCAGGGCTAGAGAGTACATAAAGTCGCTTCCCATGTACCCTGCCGCGCCACTGGAGAAGATGTTCCCTCGAGTCAACCCGAAAGGCATAGATCTTTTACAGCGTATGCTTGTTTTTGACCCTGCGAAGAGGATTACTGCTAAGGAGGCACTGGAGCATCCGTATTTGCAAACATACCACGATCCAAACGACGAACCTGAAGGCGAACCCATCCCACCCAGCTTCTTCGAGTTTGATCACTACAAGGAGGCACTAACGACGAAAGACCTCAAGAAACTCATTTGGAACGAAATATTTAGT |
| homologous long arms R' (300 bp**)** | CCATCATTATCATTAAAATATCAACCCGAAGAACAATGTATACATATACATATACGTACACATATACATATGTACATATGACATACGTATTAGCCGCTGAGGACGCGGACGTATAAAAGGACAATACTTATATGGAGCTAAGGGGAGCAGTTACGCAACTCCGTGATCGCGCGCCACGGGCCGTCGGCGGCTGTTAATTGAAGAAAAAAAAAATGAAGAACCACAAGGGGTGATCCATATAGGTGACTAGCATCATCCCCTGCGACGCGCGGCCCGCCGGGCCAAAGGCGGGCAATGCGC |
| 6aa [GS]x linker (18 bp) | ggtagcggcagcggtagc |
| Primer F (20 bp) | AAGTCGCTTCCCATGTACCC |
| Primer R (20 bP) | GTTGCGTAACTGCTCCCCTT |

Movies S1-S7: Total internal reflection microscopy images of yeast cell (*FUS3*-GFP strain) in response to 0.2 μM, 0.6 μM, 0.7 μM, 0.8 μM, 1.0 μM, 2.0 μM, and 3.0 μM pheromone over time. The shooting interval of the photos was 10 minutes/frame. Yeast cells were cultured in a microfluidic device containing the normal YNB medium for 60 minutes, and then switched to the medium containing a specific concentration of pheromone for cultivation. The green fluorescence in the cell is the gene expression level of Fus3.

Movies S8-S9: Total internal reflection microscopy images of yeast cells (*CDC24*_GFP*-FUS3*_yomCherry strain) in response to 0.8 μM pheromone over time. The shooting interval of the photos was 10 minutes/frame. Yeast cells were cultured in a microfluidic device containing the normal YNB medium for 60 minutes, and then switched to the medium containing a specific concentration of pheromone for cultivation. The green fluorescence in the cell is the gene expression level of Cdc24; the red fluorescence in the cell is the gene expression level of Fus3.

References:

1.	P. M. P. Anne-Christine Butty, Linda S. Huang, Ira Herskowitz, Matthias Peter*, The Role of Far1p in Linking the Heterotrimeric G Protein to Polarity Establishment Proteins During Yeast Mating. *SCIENCE*, (1998).

2.	D. Matheos, M. Metodiev, E. Muller, D. Stone, M. D. Rose, Pheromone-induced polarization is dependent on the Fus3p MAPK acting through the formin Bni1p. *The Journal of cell biology* **165**, 99-109 (2004).

3.	M. Peter, A. Gartner, J. Horecka, G. Ammerer, I. Herskowitz, FAR1 links the signal transduction pathway to the cell cycle machinery in yeast. *Cell* **73**, 747-760 (1993).

4.	E. Elion, B. Satterberg, J. Kranz, FUS3 phosphorylates multiple components of the mating signal transduction cascade: evidence for STE12 and FAR1. *Molecular biology of the cell* **4**, 495-510 (1993).

5.	R. L. Roberts, G. R. Fink, Elements of a single MAP kinase cascade in Saccharomyces cerevisiae mediate two developmental programs in the same cell type: mating and invasive growth. *Genes & development* **8**, 2974-2985 (1994).

6.	I. Herskowitz, MAP kinase pathways in yeast: for mating and more. *Cell* **80**, 187-197 (1995).

7.	E. A. Elion, J. A. Brill, G. R. Fink, FUS3 represses CLN1 and CLN2 and in concert with KSS1 promotes signal transduction. *Proceedings of the National Academy of Sciences* **88**, 9392-9396 (1991).

8.	E. Blackwell *et al.*, Effect of the pheromone-responsive Gα and phosphatase proteins of Saccharomyces cerevisiae on the subcellular localization of the Fus3 mitogen-activated protein kinase. *Molecular and cellular biology* **23**, 1135-1150 (2003).

9.	F. van Drogen, V. M. Stucke, G. Jorritsma, M. Peter, MAP kinase dynamics in response to pheromones in budding yeast. *Nature cell biology* **3**, 1051-1059 (2001).

10.	A. Papagiannakis, B. Niebel, E. C. Wit, M. Heinemann, Autonomous Metabolic Oscillations Robustly Gate the Early and Late Cell Cycle. *Mol Cell* **65**, 285-295 (2016).

11.	F. Caudron, Y. Barral, A super-assembly of Whi3 encodes memory of deceptive encounters by single cells during yeast courtship. *Cell* **155**, 1244-1257 (2013).

12.	A. Doncic *et al.*, Compartmentalization of a bistable switch enables memory to cross a feedback-driven transition. *Cell* **160**, 1182-1195 (2015).
